# Activation of the Keap1/Nrf2 pathway suppresses mitochondrial dysfunction in *C9orf72* ALS/FTD *in vivo* models and patient iNeurons

**DOI:** 10.1101/2023.10.02.560439

**Authors:** Wing Hei Au, Leonor Miller-Fleming, Alvaro Sanchez-Martinez, James A. K. Lee, Madeleine J. Twyning, Hiran A. Prag, Sarah Granger, Katie Roome, Laura Ferraiuolo, Heather Mortiboys, Alexander J. Whitworth

## Abstract

Mitochondrial dysfunction such as excess production of reactive oxygen species (ROS) and defective mitochondrial dynamics are common features of *C9orf72* Amyotrophic Lateral Sclerosis/Frontotemporal Dementia (ALS/FTD), but it remains unclear whether these are causative or a consequence of the pathogenic process. To address this, we have performed a comprehensive characterisation of mitochondrial dysfunction *in vivo* model, analysing multiple transgenic *Drosophila* models of *C9orf72*-related pathology, which can be correlated to disease-relevant locomotor deficits. Genetic manipulations to reverse different aspects of mitochondrial disruption revealed that only genetic upregulation of antioxidants such as mitochondrial Sod2 and catalase were able to rescue *C9orf72* locomotor deficits, suggesting a causative link between mitochondrial dysfunction, ROS and behavioural phenotypes. By analysing the Keap1/Nuclear factor erythroid 2–related factor 2 (Nrf2) pathway, a central antioxidant response pathway, we observed a blunted response in the *C9orf72* models. However, both genetic reduction of Keap1 and its pharmacological targeting by dimethyl fumarate (DMF), was able to rescue *C9orf72*-related motor deficits. In addition, analysis of *C9orf72* patient-derived iNeurons showed increased ROS that was suppressed by DMF treatment. These results indicate that mitochondrial oxidative stress is an upstream pathogenic mechanism leading to downstream mitochondrial dysfunction such as alterations in mitochondrial function and turnover. Consequently, our data support targeting the Keap1/Nrf2 signalling pathway as a viable therapeutic strategy for *C9orf72*-related ALS/FTD.

## Introduction

Amyotrophic lateral sclerosis (ALS) is characterised by the loss of upper and lower motor neurons leading to symptoms such as muscle weakness and paralysis. A plethora of evidence supports a clinical, pathologic and genetic overlap between ALS and frontotemporal dementia (FTD) (1), which is characterised by the degeneration of frontal and temporal lobes leading to clinical symptoms such as cognitive impairment and changes in behaviour and personality. Up to 50% of ALS patients report cognitive and behavioural changes; similarly, motor neuron dysfunction is a common feature in approximately 15% of FTD cases (2). A hexanucleotide repeat expansion consisting of GGGGCC (G4C2) in the first intron of *C9orf72* is the most common pathogenic mutation in ALS/FTD (3, 4). Several pathogenic mechanisms have been proposed including haploinsufficiency of the gene product and the sequestration of RNA binding proteins at accumulations (foci) of the transcribed RNA (5). However, although intronic, the expanded RNA can also be translated through a mechanism known as repeat associated non-AUG (RAN) translation which produces 5 different dipeptide repeat proteins (DPRs), with arginine DPRs exhibiting the most toxicity (6–14).

Abundant evidence supports that mitochondrial dysfunction is an early alteration in ALS (15, 16), and mitochondrial bioenergetics and morphological changes have been observed in *C9orf72* patient fibroblasts and iPSC-derived motor neurons (17–19). Cellular and mouse models expressing poly-GR have consistently shown mitochondrial perturbations such as redox imbalance and DNA damage (20). Evidence indicates that poly-GR binds to ATP5A1 thereby compromising mitochondrial function (21), and that poly-GR toxicity and aggregation can also occur due to frequent stalling of poly-GR translation on the mitochondrial surface, triggering ribosome-associated quality control and C-terminal extension (22).

Various cellular mechanisms exist to counteract disruptions in mitochondrial function and redox imbalance, from antioxidant defence mechanisms to the wholesale degradation of mitochondria via macroautophagy (mitophagy), which has been relatively understudied in the *C9orf72* context. One of the most important upstream mechanisms that counteract redox imbalance is the Keap1/Nuclear factor erythroid 2–related factor 2 (Nrf2) pathway. Under basal conditions, Nrf2, a master regulatory transcription factor for antioxidant and cell-protective factors, is negatively regulated via targeted proteasomal degradation by an E3 ubiquitin ligase adaptor, Keap1. This interaction is alleviated upon oxidative and electrophilic stresses, and can also be provoked by Keap1 small-molecule inhibitors, causing Nrf2 to accumulate in the nucleus. This induces the expression of a repertoire of protective factors such as proteins with detoxification, antioxidant and anti-inflammatory properties, with the purpose to maintain mitochondrial function and redox balance (23, 24).

In this study, we have conducted an extensive *in vivo* characterisation of mitochondrial dysfunction in multiple *Drosophila* models of *C9orf72* ALS/FTD by studying mitochondrial dynamics, respiration, mitophagy and redox homeostasis, in conditions that cause the disease-relevant locomotor deficits. We found that only reversal of oxidative stress by the overexpression of antioxidant genes was able to rescue the progressive loss of motor function. Focussing on a role for the key antioxidant Keap1/Nrf2 signalling pathway, we sought to investigate genetic and pharmacological inactivation of the negative regulator Keap1 in the *C9orf72 Drosophila* models as well as *C9orf72* patient-derived iNeurons. Our results suggest that mitochondrial oxidative stress is an upstream pathogenic mechanism and activation of the Keap1/Nrf2 pathway could be a viable therapeutic strategy for ALS/FTD.

## Methods

### *Drosophila* stocks and husbandry

Flies were raised under standard conditions in a temperature-controlled 12:12 hour light/dark cycle incubator at 25 °C and 65% relative humidity, on standard *Drosophila* food containing cornmeal, agar, molasses, yeast and propionic acid (Genetics Department, University of Cambridge). Transgene expression was induced using the pan-neuronal driver *nSyb-*GAL4 or the predominantly motor neuron driver *DIPγ-*GAL4. The following strains were obtained from Bloomington *Drosophila* Stock Center: *w*^1118^ (RRID:BDSC_6326), *nSyb-* GAL4 (RRID:BDSC_51635), *UAS-mito-HA-GFP* (RRID:BDSC_8443), *UAS-luciferase RNAi* (RRID:BDSC_31603), *UAS-Sod1* (RRID:BDSC_33605), *UAS-Sod2* (RRID:BDSC_24492), *UAS-catalase* (RRID:BDSC_24621), *Opa1^s3475^* (RRID:BDSC_12188), *UAS-GFP.mCherry.Atg8a* (RRID:BDSC_37749), *UAS-mito-roGFP2-Orp1* (RRID:BDSC_67667), *UAS-cyto-Grx1-roGFP2* (RRID:BDSC_67662), *UAS-mito-roGFP2-Grx1* (RRID:BDSC_67664); the Zurich ORFeome Project: *UAS-LacZ* (F005035) and *UAS-Tango11.HA* (F002828); and the Fly Stocks of National Institute of Genetics: *UAS-USP30 RNAi* (3016R-2). Other *Drosophila* lines were kindly provided as follows: *DIPγ* [MI03222]*-* GAL4 (RRID:BDSC_90315) from Dr Robert Carrillo (25) (26), *UAS-G4C2×3*, *UAS-G4C2×36* and *UAS-GR36* from Prof. Adrian Isaacs (6), *UAS-GR1000-eGFP* from Dr Ryan West (27), *UAS-mito.Catalase* from Prof. Alberto Sanz (28), *UAS-Drp1.WT* from Prof. Jongkyeong Chung (29), *Marf^B^* from Prof. Hugo Bellen (30), *GstD1-GFP* (31) and *Keap1^del^* (32) from Prof. Linda Partridge. The *UAS-mito-QC* reporter (RRID:BDSC_91640) has been described before (33).

### Drug treatments

For paraquat treatment, standard *Drosophila* food was supplemented with paraquat (Sigma, 856177) to a final concentration of 10 mM. For dimethyl fumarate (DMF, Sigma, 242926) adult treatment, DMF (or an equivalent volume of ethanol for the vehicle control) was added into a sugar-yeast (SY) medium (32) consisting of 15 g/L agar, 50 g/L sugar and 100 g/L yeast to a final concentration of 7 μM. All flies were transferred into freshly prepared supplemented food every 2-3 days. For larval DMF treatment, crosses of the parental genotypes were set up in standard food containing 1 μM DMF (or an equivalent volume of ethanol for the vehicle control), providing the larval offspring exposure to the treatment throughout development.

### Locomotor and survival assays

#### Larval crawling

Larval crawling was conducted by using wandering third instar (L3) larvae. Each larva was placed in the middle of a 1 % agar plate, where they were left to acclimatise for 30 seconds. After, the number of forward and backward peristaltic waves were counted for 60 seconds and recorded.

#### Climbing

The repetitive iteration startle-induced negative geotaxis (RISING, or ‘climbing’) assay was performed using a counter-current apparatus as previously described (Greene et al., 2003). Briefly, groups of 15-22 flies were placed in a temperature-controlled room for 30 minutes for temperature acclimatisation and transferred to test tubes for another 30 minutes. Flies were placed into the first chamber, tapped to the bottom, and given 10 seconds to climb a 10 cm distance. 20-day-old flies were given 20 seconds to climb to account for overall reduced mobility. Flies that reached the upper portion, i.e., climbed 10 cm or more, were shifted into the adjacent chamber. After five successful trials, the number of flies in each chamber was counted and the average score was calculated and expressed as a climbing index.

#### Survival

For survival lifespan experiments, groups of 20 males each were collected with minimal time (<30 s) under light anaesthesia and placed in separate food vials in standard food and transferred every 2-3 days to fresh food and the number of dead flies recorded. Percent survival was calculated at the end of the experiment using https://flies.shinyapps.io/Rflies/ (Luis Gracia).

#### Oxidative stress assays

For short term hydrogen peroxide (H_2_O_2_) and paraquat (PQ) assays, flies were kept in vials containing filter paper soaked in 5% w/v sucrose solution containing either 1 % H_2_O_2_ (Fisher Scientific, 10386643) or 10 mM PQ (Sigma, 856177). Deaths were recorded two times per day, and flies were transferred every 2-3 days to fresh tubes.

### Immunohistochemistry and sample preparation

For mitophagy analysis with mito-QC reporter, autophagy analysis with GFP.mCherry.Atg8a reporter and mitochondrial morphology analysis, third instar larval brains were dissected in PBS and fixed in 4% formaldehyde (FA) (Thermo Scientific, 28908) in phosphate buffered saline (PBS) for 20 minutes at room temperature (RT). For the mitophagy and autophagy experiments, the 4 % FA/PBS was adjusted to pH 7. Samples were then washed in PBS followed by water to remove salts. ProLong Diamond Antifade mounting medium (Invitrogen, P36961) was used to mount the samples and imaged the next day.

For immunostaining of third instar larval brains, larvae were dissected and fixed as described and permeabilised in 0.3 % Triton X-100 in PBS (PBS-T) for 30 minutes, blocked with 1% bovine serum albumin (BSA) in PBS-T for 1 hour at RT. Tissues were then incubated with rabbit anti-CncC (kind gift from Dr Fengwei Yu), diluted in 1% BSA in PBS-T overnight at 4 °C, then washed 3 times 10 min with PBS-T, followed by incubation with secondary antibody – goat anti-rabbit IgG H&L Alexa Fluor™ 488 (1:500, Invitrogen, A11008). The tissues were washed in PBS-T with 1:10,000 Hoechst (Invitrogen, H3570) for 20 mins followed by 2 times PBS washes and mounted on slides using ProLong Diamond Antifade mounting medium.

For immunostaining of adult brains, flies were dissected in PBS and fixed on ice in 4 % FA in PBS for 30 minutes. Brains were washed three times for 20 minutes in PBS-T, prior to blocking for 4 hrs in 4% normal goat serum (NGS) in PBS-T. Tissues were then incubated with rabbit anti-CncC (kind gift from Dr Fengwei Yu), diluted in 4 % NGS in PBS-T overnight at 4 °C, then washed 3 times 10 min with PBS-T, followed by incubation with secondary antibody for 2 hours. The tissues were washed in PBS-T with 1:10,000 Hoechst for 20 mins followed by 2 times PBS washes and mounted on slides using Prolong Diamond Antifade mounting medium.

### ROS analysis

#### MitoSOX

2-3 10-day-old adult brains were dissected at a time in PBS, incubated in 20 μM of MitoSOX Red (Invitrogen, M36008) for 30 minutes in the dark, washed with PBS for three times, mounted on poly-L-lysine coated wells on a 1.5 mm coverslip and imaged live immediately. The maximum intensity of projected z stacks from imaged brains was quantified using ImageJ.

#### Dihydroethidium (DHE)

2-3 larval brains were dissected in HL3 (5 mM HEPES, 70 mM NaCl, 5 mM KCl, 20 mM MgCl_2_, 1.5 mM CaCl_2_, 5 mM Trehalose, 115 mM Sucrose) (34) and incubated for 15 minutes in the dark at RT in 30 μM of DHE (Invitrogen, D11347). Samples were then washed in HL3 for 10 minutes before imaging live in a drop of HL3 on poly-L-lysine coated wells on 1.5 mm coverslips.

#### Genetic roGFP2 reporters

Mitochondrial and cytosolic ROS imaging was performed using the *mito-roGFP2-Orp1, mito-roGFP2-Grx1* and *cyto-Grx1-roGFP2* reporter lines. Third larval instar brains were dissected in HL3 and placed in a drop of HL3 on poly-L-lysine coated wells on 1.5mm coverslip and imaged by excitation at 488 nm (reduced) or 405 nm (oxidized), with emission detected at 500-530 nm. The maximum intensity of projected z-stacks from imaged brains was quantified using FIJI (Image J) and the ratio of 405/488 nm was calculated.

### Microscopy

Fluorescence imaging was conducted using a Zeiss LSM 880 confocal microscope (Carl Zeiss MicroImaging) equipped with Nikon Plan-Apochromat 63x/1.4 NA oil immersion objectives. Images were prepared using FIJI (Image J). For mito-QC imaging, the Andor Dragonfly spinning disk microscope was used, equipped with a Nikon Plan-Apochromat 100x/1.45 NA oil immersion objective and iXon camera. Z-stacks were acquired with 0.2 μm steps. For larval morphology, images were acquired using a Leica DFC490 camera mounted on a Leica MZ6 stereomicroscope.

### Quantification and analysis methods

#### Mitophagy

Confocal images were processed using FIJI (Image J). The quantification of mitolysosomes was performed as described in (33) using Imaris (version 9.0.2) analysis software. Briefly, a rendered 3D surface was generated corresponding to the mitochondrial network (GFP only). This surface was subtracted from the mCherry signal which overlapped with the GFP-labelled mitochondrial network, defining the red-only mitolysosomes puncta with an estimated size of 0.5 μm and a minimum size cut-off of 0.2 μm diameter determined by Imaris.

#### Autophagy

The quantification of autolysosomes was performed using FIJI (Image J) with the 3D Objects Counter Plugin. An area of interest was selected by choosing 6-10 cells per image. The threshold was based on matching the mask with the fluorescence. A minimum size threshold of 0.05 μm^3^ was set to select autolysosomes.

#### Mitochondrial morphology

After acquisition of images, each cell was classified using a scoring system where morphology was scored as fragmented, WT/tubular or fused/hyperfused. All images were blinded and quantified by three independent investigators. Data presented in Figure 3 and Supplementary Figure 2 were conducted concurrently, therefore the control groups are the same but replicated in the two figures for ease of reference.

#### Nuclear CncC quantification

All acquired images were taken with the same laser and gain during acquisition, which allowed a threshold to be set in FIJI (Image J) that was consistent for all images. For each brain, using the Hoechst signal, 10 nuclei from the central part of the larval CNS and 10 from the periphery were quantified to minimise bias. This was overlaid onto the CncC channel and the mean intensity within the nuclei was measured using the ROI manager. All images were blinded before quantification.

### Mitochondrial respiration

Mitochondrial respiration was monitored at 25°C using an Oxygraph-2k high resolution respirometer (OROBOROS Instruments). Standard oxygen calibration was performed before the start of every experiment. Twenty 5-day-old adult fly heads per replicate for each genotype was extracted using forceps and placed in 100 μL of Respiration buffer (RB) (120 mM sucrose, 50 mM KCl, 20 mM Tris-HCl, 4 mM KH_2_PO4, 2 mM MgCl_2_, 1 mM EGTA and 1 g/L fatty acid-free BSA, pH 7.2). This was homogenised on ice using a pestle with 20 strokes. 1 mL of RB was added to the homogenate and passed through a 1 mL syringe with a piece of cotton wool inside to remove the debris. This was repeated with another 1 mL of RB. In total, 2.1 mL of homogenate was added into the respiratory chambers. For coupled ‘state 3’ assays, saturating concentrations of substrates including 10 mM glutamate, 2 mM malate, 10 mM proline and 2.5 mM ADP was added to measure Complex I-linked respiration. 0.15 μM rotenone was added to inhibit Complex I and 10 mM succinate was added to measure Complex II-linked respiration. Data acquisition and analysis were carried out using Datlab software (OROBOROS Instruments).

### Immunoblotting

Proteins were isolated from either larval brains or adult heads using RIPA lysis buffer (50 mM pH 7.4 Tris, 1 M NaCl, 0.1 % SDS, 0.5 % sodium deoxycholate and 1 % NP-40) supplemented with cOmplete mini EDTA-free protease inhibitors (Roche). After protein quantification using bicinchoninic acid assay (BCA) (Thermo Scientific, 23225), Laemmli buffer (BioRad, 1610747) containing 1:10 β-mercaptoethanol (Sigma, M6250) was added. Samples were boiled at 95 ℃ for 10 minutes and resolved by SDS-PAGE using 4-20 % or 10 % precast gels (BioRad, 4561093), depending on the molecular weight of the desired protein, and transferred onto nitrocellulose membrane (BioRad, 1704158) using a semi-dry BioRad TransBlot system. Membranes were blocked with 5 % (w/v) dried skimmed milk powder (Marvel Instant Milk) in Tris-buffered saline (TBS) with 0.1 % Tween-20 (TBS-T) for 1 hour at RT and probed with the appropriate primary antibodies diluted in TBS-T overnight at 4 ℃. After three 10-minute washes in TBS-T, the membranes were incubated with the appropriate horse radish peroxidase (HRP-conjugated) secondary antibodies diluted in 5 % milk in TBS-T for 1 hour at RT. Membranes were washed three times for 10 minutes in TBS-T and detection was achieved with Amersham ECL-Prime detection kit (Cytiva RPN2232). Blots were imaged using the Amersham Imager 680 with further analysis and processing using FIJI (Image J).

The following primary antibodies for immunoblotting were used in this study: GFP (1:100, Abcam, ab294), tubulin (1:5000, Sigma, T9026), ref(2)P (1:1000, Abcam, ab178440) and GABARAP (1:1000, Abcam, ab109364). The following secondary antibodies for immunoblotting were: goat anti-mouse IgG H&L (Horse radish peroxidase (HRP) conjugated, 1:10,000, Abcam, ab6789) and goat anti-rabbit IgG H&L HRP (1:10,000, Thermo Fisher Scientific, G21234).

### qRT-PCR

Quantitative real-time PCR (qRT-PCR) was carried out as follows: 30 heads were collected and placed in a 2 mL tube containing 1.4 mm ceramic beads (Fisherbrand, 15555799) for tissue preparation. 400 μL of TRI Reagent (Sigma, T9424) was added and placed into Minilys homogeniser (Bertin Technologies) where the programme was set to maximum speed for 10 seconds. The samples were placed back on ice for 5 minutes before two further rounds of lysis. Direct-zol™ RNA MiniPrep kit (Zymo Research, R2050) was used to extract RNA following manufacturer’s instructions. TURBO DNA-free™ Kit (Invitrogen, AM1907) was used to remove contaminating DNA by following manufacturer’s instructions. cDNA synthesis was achieved by using Maxima™ H Minus cDNA Synthesis Kit (Thermo Scientific, M1681) following manufacturer’s instructions. Equivalent (500 μg) total RNA underwent reverse transcription for each sample. Finally, qRT-PCR was ran using Maxima SYBR Green/ROX Kit (Thermo Scientific, K0221) following manufacturer’s instructions using the Quant Studio 3 RT-PCR machine. The relative transcript levels of each target gene were normalised to a geometric mean of *RpL32* and *Tub84b* reference genes; relative quantification was performed using the comparative C_T_ method entering into account PCR primer efficiency (35).

The following primers were used in this study:

*Sod1*: CCAAGGGCACGGTTTTCTTC, CCTCACCGGAGACCTTCAC
*Sod2*: GTGGCCCGTAAAATTTCGCAAA, GCTTCGGTAGGGTGTGCTT
*Catalase*: CCAAGGGAGCTGGTGCTT, ACGCCATCCTCAGTGTAGAA
*RpL32*: AAACGCGGTTCTGCATGAG, GCCGCTTCAAGGGACAGTATCTG
*Tub84b*: TGGGCCCGTCTGGACCACAA, TCGCCGTCACCGGAGTCCAT

### iNPC tissue culture, iNeuron differentiation and compound treatment

Induced Neuronal Progenitor Cells (iNPCs) were generated as previously described (36, 37). Details for the patient lines used in this study are provided in Table 1. iNPCs were maintained in DMEM/F-12 with GlutaMAX (Gibco, 31331028) supplemented with 1% N-2 (Invitrogen, 17502001), 1% B-27 (Invitrogen, 17504001) and 20 ng/ml FGF-basic (PeproTech, 100-18B) on fibronectin (Millipore, FC010) coated cell culture dishes. Cells were routinely subcultured every 2-3 days using Accutase (Corning, 15323609) to detach them. To achieve neuronal differentiation, iNPCs were plated into 6 well plates and cultured for 2 days in DMEM/F-12 media with GlutaMAX supplemented with 1% N-2, 2% B-27, and 0.5 μM DAPT (Sigma, D5942). On day 3, DAPT was removed, and media was supplemented with 0.5 μM Smoothened Agonist (SAG, Millipore, 566660), 2.5 μM Forskolin (Cayman Chemical, 11018) and 1 μM retinoic acid (Sigma, R2625) for 16 days. Cells were treated with edaravone (EDV, Sigma, M70800) and dimethyl fumarate (DMF, Sigma, 242926) for 24 hours prior to imaging. Cells were treated with 30 μM or 100 μM EDV, or DMF at concentrations of 3 μM or 10μM.

**Table 1.**
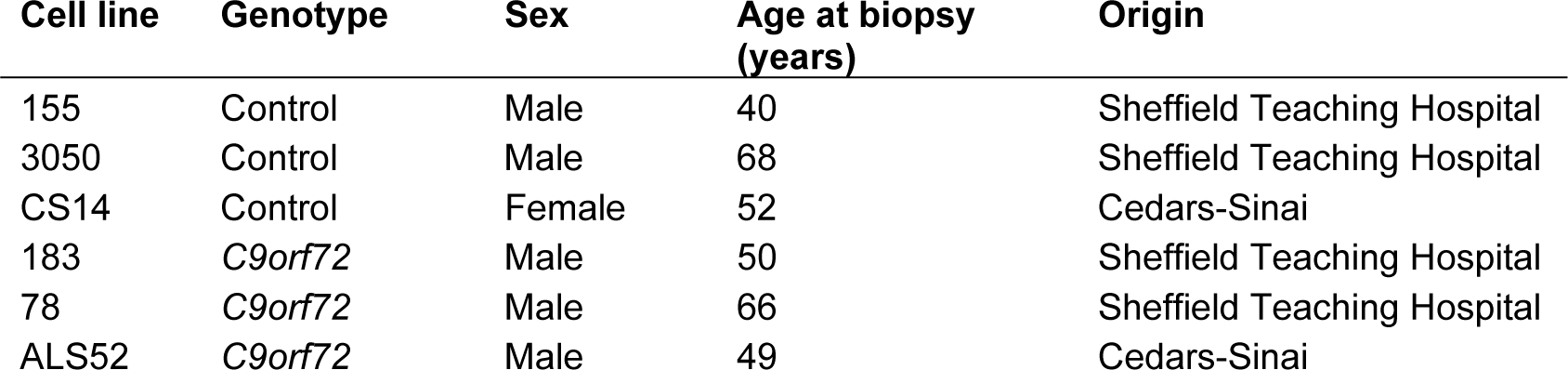
Cell lines used in this study.

| Cell line | Genotype | Sex | Age at biopsy (years) | Origin |
| --- | --- | --- | --- | --- |
| 155 | Control | Male | 40 | Sheffield Teaching Hospital |
| 3050 | Control | Male | 68 | Sheffield Teaching Hospital |
| CS14 | Control | Female | 52 | Cedars-Sinai |
| 183 | *C9orf72* | Male | 50 | Sheffield Teaching Hospital |
| 78 | *C9orf72* | Male | 66 | Sheffield Teaching Hospital |
| ALS52 | *C9orf72* | Male | 49 | Cedars-Sinai |

### Live cell imaging assays of iNeurons

Cells were stained for 30 minutes with Hoechst (Sigma, B2883) and MitoSOX Red (Invitrogen, M36008) at concentrations of 20 μM and 500 nM, respectively. Cells were imaged using an Opera Phoenix high content imaging system, with analysis performed using a custom protocol on Harmony software (PerkinElmer).

### Fixed cell imaging of iNeurons

On day 18 of differentiation, cells were fixed in 4% paraformaldehyde for 30 minutes. After PBS washes, cells were permeabilised with 0.1% Triton X-100 for 10 minutes and blocked with 5% horse serum (Sigma, H0146) for 1 hour. Cells were incubated with primary NRF2 antibody (1:1000, Abcam, ab31163) overnight at 4 ℃. Cells were washed with 0.1% Tween in PBS and incubated with donkey anti-rabbit IgG H&L Alexa Fluor™ 568 secondary antibody (1:1000, Invitrogen, A10042) and 1 μM Hoechst prior to imaging. Cells were imaged using an Opera Phoenix high content imaging system, with analysis performed using a custom protocol on Harmony software (PerkinElmer).

### Statistical analysis

GraphPad Prism 9 (RRID:SCR_002798) was used to perform all statistical analyses. Climbing data was analysed using Kruskal-Wallis non-parametric test with Dunn’s correction for multiple comparisons. Data are presented as mean ± 95% confidence interval (CI). Quantifications of larval crawling, number of mitolysosomes and autolysosomes, WB and qRT-PCR were analysed using one-way ANOVA with Bonferroni post hoc test for multiple comparisons. Data are presented as mean ± SD. Mitochondria morphology was quantified using Chi squared test. Unpaired t-tests with Welch’s correction for unequal standard deviation was used for respiratory analysis and western blots. Two-way ANOVA with Dunnett’s multiple comparison test was used for analysis of the iNeurons work. Lifespan experiments were analysed using the Mantel-Cox log-rank test. All statistical tests and n numbers are stated in the figure legends.

## Results

### *C9orf72 Drosophila* models present a wide range of mitochondrial defects

Mitochondrial dysfunction has been suggested to be directly involved in disease pathogenesis in ALS/FTD but evidence from animal models is limited (38). To comprehensively assess different aspects of mitochondrial dysfunction in *C9orf72*-related pathology *in vivo*, we utilised several previously established *Drosophila* models based on the inducible GAL4-UAS expression system (39). Expression of 36x G4C2 repeats (G4C2×36) and 36-repeat glycine-arginine DPR (GR36) are substantially neurotoxic compared to the 3x G4C2 repeats (G4C2×3) control (6). Indeed, we found that strong pan-neuronal expression of G4C2×36 (via *nSyb-* GAL4) perturbed development, where larvae were morphologically thinner compared to control (Fig. 1A), and with significantly reduced motor ability assessed by measuring larval crawling behaviour (Fig. 1B). Consistent with previous reports, pan-neuronal expression of GR36 caused even stronger phenotypes (Fig. 1A, B), and was developmental lethal at the third-instar (L3) larval stage.

**Figure 1.**
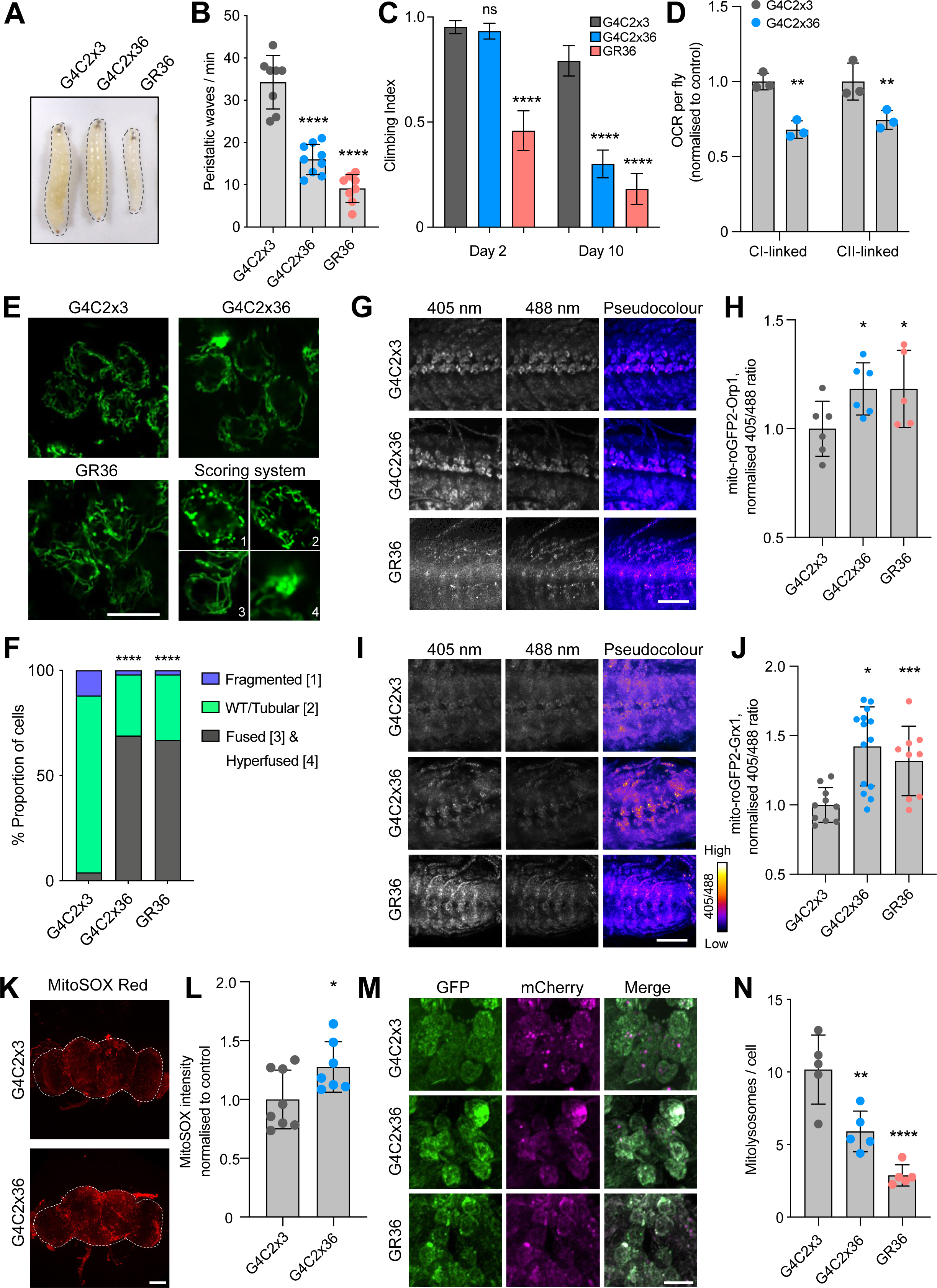
Multiple aspects of mitochondrial function are disrupted in *Drosophila C9orf72* models. **(A)** Morphology and **(B)** crawling ability of larvae expressing G4C2×3, G4C2×36 or GR36 via pan-neuronal driver *nSyb-*GAL4 (mean ± SD; one-way ANOVA with Bonferroni’s multiple comparison test; ****p < 0.0001; n = 8-10 larvae). **(C)** Climbing analysis of 2- and 10-day-old adults expressing G4C2×3, G4C2×36 and GR36 flies via predominantly motor neuron driver *DIPγ-*GAL4 (mean ± 95% CI; Kruskal-Wallis non-parametric test with Dunn’s correction; ****p < 0.0001; n = 55-100 flies). **(D)** Oxygen consumption rate (OCR) of 5-day-old fly brains of G4C2×3 and G4C2×36 using *nSyb*-GAL4 (mean ± SD; paired t-test with Welch’s corrections; **p < 0.01; n=3 biological replicates). **(E)** Confocal microscopy of larval neurons where mitochondria are labelled with pan-neuronal expression of mito.GFP, co-expressing G4C2×3, G4C2×36 or GR36 with *nSyb*-GAL4. Scale bar = 10 µm. **(F)** Quantification of **E** on a cell-by-cell basis as [1] fragmented, [2] tubular (WT) appearance, [3] fused, or [4] hyperfused (chi-squared test; ****p < 0.0001; n = 8-10 larvae). **(G -J)** Confocal microscopy of **(G)** mito-roGFP2-Orp1 mitochondrial H_2_O_2_ reporter and **(I)** mito-roGFP2-Grx1 mitochondrial glutathione redox potential reporter co-expressed with G4C2×3, G4C2×36 and GR36 in the larval ventral ganglion with *nSyb*-GAL4. Representative pseudocolour ratio images are depicted and **(H, J)** the 405/488 ratio quantified. Scale bar = 50 µm (mean ± SD; one-way ANOVA with Bonferroni’s multiple comparison test; *p < 0.05, ***p < 0.001; n = 5-14 larvae). **(K, L)** MitoSOX staining in 5-day-old G4C2×3 and G4C2×36 brains (pan-neuronal expression with *nSyb*-GAL4). White dotted line indicates brain boundaries. Scale bar = 100 µm (mean ± SD; unpaired t-test with Welch’s corrections; *p < 0.05). **(M, N)** Confocal analysis of the mito-QC mitophagy reporter co-expressed with G4C2×3, G4C2×36 and GR36 in the larval ventral ganglion with *nSyb*-GAL4. Scale bar = 10 µm (mean ± SD; one-way ANOVA with Bonferroni’s multiple comparison test; **p < 0.01, ****p < 0.0001; n = 5 larvae).

In order to analyse *C9orf72* pathology in the adult stage, where the impact of ageing can also be investigated, we found that expression of the *C9orf72* transgenes via a predominantly motor neuron driver, *DIPγ-*GAL4, was adult-viable and caused an age-related decline in motor ability assessed using the negative geotaxis ‘climbing’ assay (Fig. 1C). While motor neuron expression of G4C2×36 did not impact motor behaviour at 2-days-old, by 10 days, G4C2×36 flies exhibited significantly impaired climbing ability (Fig. 1C). As before, motor neuron expression of GR36 was more toxic, causing a significant motor deficit at 2-days-old which worsened by 10 days (Fig. 1C). A similar pattern was observed in the lifespan of *DIPγ-*GAL4 driven G4C2×36 and GR36 flies with a mild phenotype from G4C2×36 but a markedly shortened lifespan with GR36 (Fig. S1A).

Previous studies analysing mitochondrial defects in *Drosophila C9orf72* models have mostly focussed on muscle-directed expression of poly-GR (22, 40). Here we aimed to explore the potential involvement of mitochondrial dysfunction in a neuronal context. Initially, we observed that pan-neuronal expression of G4C2×36 caused a significant reduction in complex I- and complex II-linked respiration in lysates from young (5-day-old) fly heads (Fig. 1D), indicative of a generalised mitochondrial disruption. To probe this in more detail, we next analysed mitochondrial morphology in larval neurons upon pan-neuronal expression of the *C9orf72* transgenes. As mitochondria are dynamic organelles, microscopy analysis of mito.GFP-labelled mitochondria typically reveals a mix of short, round (fragmented) and long, tubular (fused) morphologies (Fig. 1E), which can be quantified using a scoring system that characterises the overall mitochondrial morphology on a cell-by-cell basis (Fig. 1E, F). Using this approach, we found that mitochondria were more elongated and hyperfused in G4C2×36 and GR36 expressing neurons (Fig. 1E, F).

Mitochondrial morphology is known to respond to changes in reactive oxygen species (ROS) levels as well as other physiological stimuli. First, we tested the susceptibility of G4C2×36 flies to oxidative stress induced by paraquat (generates superoxide anions) and hydrogen peroxide (H_2_O_2_ - generates hydroxyl radicals). We observed that G4C2×36 flies were hypersensitive to both types of oxidative stressors when compared to control animals (Fig. S1B, C).

ROS can occur in different forms (e.g., superoxide anions, H_2_O_2_) which may vary in their prevalence and functional significance across cellular compartments (mitochondrial vs cytosolic). Therefore, it is important to study the effects of ROS species in each compartment to understand the tight regulation required to achieve redox homeostasis. We utilised genetically encoded redox-sensitive fluorescent protein (roGFP2) probes where fusion of roGFP2 to oxidant receptor peroxidase 1 (Orp1) or glutaredoxin 1 (Grx1) results in roGFP2 oxidation by H_2_O_2_ or oxidised glutathione (GSSG), respectively, which leads to a shift in the excitation maxima of the fusion constructs from 488 to 405 (41). Importantly, these reporters can be targeted to specific subcellular compartments, i.e., cytosol or mitochondria, giving insights into compartment-specific redox status. We observed a significant increase in the oxidised status of mitochondrial roGFP2-Orp1 (Fig. 1G, H) and mitochondrial roGFP2-Grx1 (Fig. 1I, J) in larval neurons when combined with G4C2×36 and GR36, indicating an increase in mitochondrial H_2_O_2_ and GSSG in these animals. In contrast, no significant differences were observed in cytosolic glutathione redox potential (E_GSH_) (Fig. S1D), and little or no changes were detected with dihydroethidium (DHE) intensity, an indicator of cytosolic superoxide, in G4C2×36 and GR36 larvae (Fig. S1E). To extend our analysis of the larval model to adult flies, and to investigate an independent measure of ROS, we used MitoSOX to investigate levels of mitochondrial superoxide in the adult brain. In agreement with the fluorescent reporter results, we observed an increase in MitoSOX intensity in 5-day-old G4C2×36 flies compared to G4C2×3 control (Fig. 1K, L). Taken together, these data suggest that mitochondrial but not cytosolic redox state is disrupted in these *Drosophila* models of *C9orf72*-related pathology. These results are summarised in Table 2.

**Table 2.**
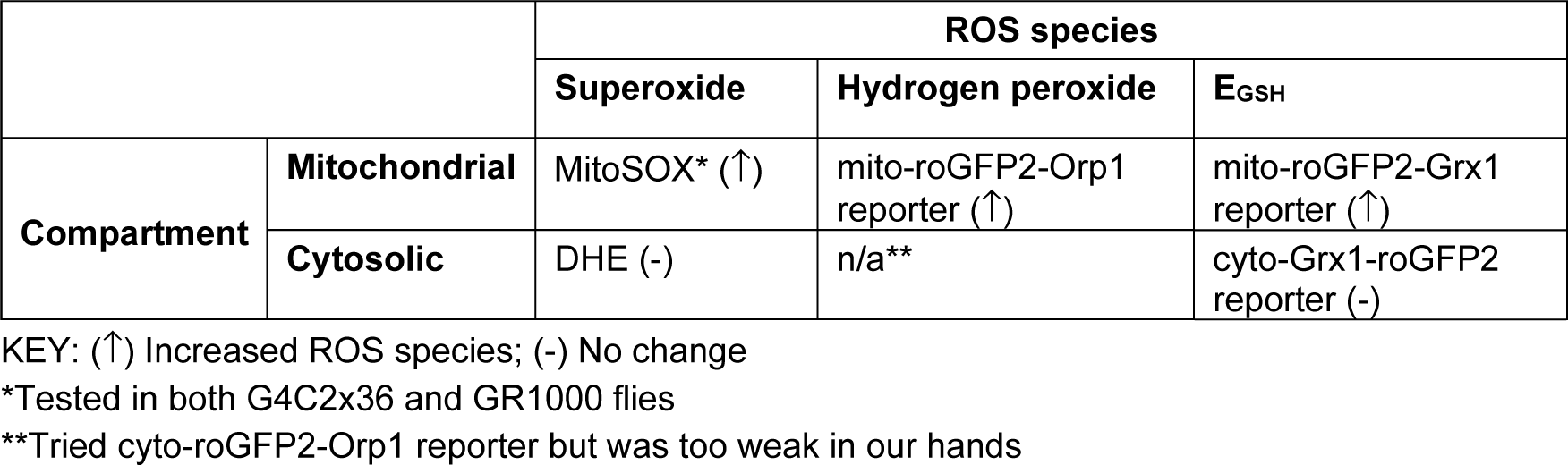
Summary of ROS 418 outcomes in G4C2×36 flies.

|  |  | ROS species |  |  |
| --- | --- | --- | --- | --- |
|  |  | Superoxide | Hydrogen peroxide | E<sub>GSH</sub> |
| Compartment | Mitochondrial | MitoSOX* (↑) | mito-roGFP2-Orp1 reporter (↑) | mito-roGFP2-Grx1 reporter (↑) |
|  | Cytosolic | DHE (-) | n/a** | cyto-Grx1-roGFP2 reporter (-) |

KEY: (↑) Increased ROS species; (-) No change
\*Tested in both G4C2x36 and GR1000 flies
\*\*Tried cyto-roGFP2-Orp1 reporter but was too weak in our hands

Changes in mitochondrial morphology and increases in ROS can be indicative of ongoing mitochondrial damage which can be alleviated via mitophagy. Thus, we next used the mito-QC mitophagy reporter (33) to investigate the mitophagy status in the *C9orf72* models. Briefly, the mito-QC reporter uses a tandem GFP-mCherry fusion protein targeted to the outer mitochondrial membrane (OMM). Mitolysosomes are marked when GFP is quenched by the acidic environment but mCherry fluorescence is retained, resulting in ‘red-only’ puncta (33). Analysis showed that there were significantly fewer mitolysosomes in G4C2×36 and GR36 larval neurons compared to G4C2×3 control (Fig. 1M, N), suggesting that mitophagy is perturbed. However, this was not a mitophagy-specific defect as the analogous reporter for general autophagy, GFP-mCherry-Atg8a autophagy reporter (fly Atg8a is homologous to LC3), also showed a reduction in the number of mCherry-positive autolysosomes in G4C2×36 and GR36 larval neurons (Fig. S1F). Consistent with this, immunoblot analysis of protein lysates from 5-day-old G4C2×36 fly heads revealed an increase in ref(2)P levels (fly homologue of p62) as well as a reduction in lipidated Atg8a-II levels (Fig. S1G, H). These results are consistent with a reduction in general autophagic flux, which could impact mitophagic flux, in agreement with previous literature using different *C9orf72* models (42).

Recently, West et al. (27) developed additional *C9orf72* models which express physiologically relevant *C9orf72*-related DPRs, i.e., ∼1000 repeats. For comparison to the Mizielinska et al. lines, we also analysed the GR1000-eGFP line (for simplicity, hereafter called GR1000). We observed an age-related decline in motor performance compared to control with pan-neuronal expression of GR1000, where GR1000 flies were not able to climb at all by 20 days (Fig. 2A), similar to that previously reported (27). To complement our results shown in the G4C2×36 and GR36, we measured mitochondrial superoxide with MitoSOX staining in GR1000 adult brains as it was not possible to use the roGFP2 reporters in conjunction with eGFP-tagged GR1000. Here, we also observed an increase in MitoSOX staining in 10-day-old GR1000 brains compared to control (Fig. 2B, C).

**Figure 2.**
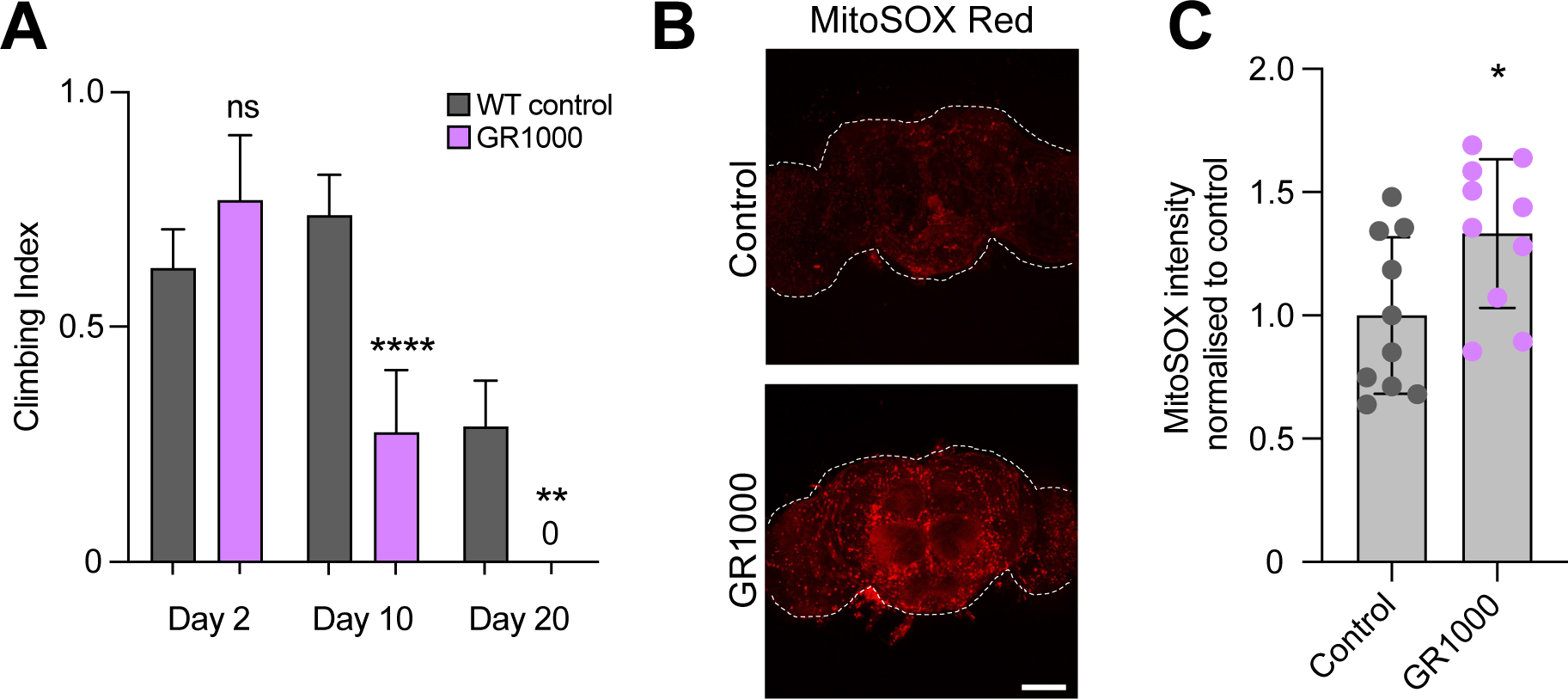
GR1000 flies show increased mitochondrial ROS. **(A)** Climbing analysis of 2-10- and 20-day-old adults of pan-neuronal expression, via *nSyb-* GAL4, of mito.GFP (WT control) or GR1000 (mean ± 95% CI; Kruskal-Wallis non-parametric test with Dunn’s correction; ns=non-significant, **p < 0.01, ****p < 0.0001; n = 60-100 flies). **(B, C)** Confocal microscopy analysis of MitoSOX staining in 10-day-old adult brains in WT control (*nSyb>mito.GFP*) and GR1000 flies (pan-neuronal expression with *nSyb*-GAL4). White dotted line indicates adult brain boundaries. Scale bar = 100 µm (mean ± SD; unpaired t-test with Welch’s corrections; *p < 0.05 n = 10 brains).

Taken together, the preceding data show that various aspects of mitochondrial form and function are disturbed in multiple *Drosophila* models of *C9orf72* pathology which correlate with the organismal decline and loss of motor function.

### Genetic manipulation of mitochondrial dynamics or mitophagy does not rescue *C9orf72* phenotypes

We next sought to determine whether disruptions to these observed changes in mitochondrial morphology, defective mitophagy and oxidative stress are contributing factors to the neurodegenerative process. First, we evaluated whether genetic manipulations to counteract the elongated, hyperfused mitochondrial morphology caused by G4C2×36 and GR36 expression may rescue the larval locomotor defect. Thus, we combined pan-neuronal expression of G4C2×36 and GR36 with genetic manipulations to promote fission (overexpression of pro-fission factors *Drp1* or *Mff* (*Tango11* in *Drosophila*)) or reduce fusion (loss of pro-fusion factors *Opa1* and *Mitofusin* (*Marf* in *Drosophila*)). Although the elongated mitochondria observed in the G4C2×36 and GR36 neurons was partially reversed by these manipulations as expected (Fig. S2A, B), this did not result in any improvement in larval locomotion (Fig. S2C). These data suggest that excess mitochondrial fusion observed in the *C9orf72* models does not play a key role in *C9orf72* pathogenesis and may be a downstream consequence.

Similarly, since we observed reduced mitophagy in the *C9orf72* models, we aimed to reverse this by boosting mitophagy by targeting the mitophagy inhibitor USP30. Knockdown of *USP30* has been shown to rescue defective mitophagy caused by pathogenic mutations in *PRKN* and improve mitochondrial integrity in *parkin*- or *Pink1*-deficient flies (43). We have also seen that *USP30* knockdown increases mitolysosome number in *Drosophila* neurons and muscle (44), indicative of increased mitophagy. As expected, we observed that *USP30* knockdown significantly increased the number of mitolysosomes in control larval neurons (Fig. S2D, E). However, when co-expressed with G4C2×36 and GR36, *USP30* knockdown was not sufficient to rescue the reduced mitophagy (Fig. S2D, E). Moreover, pan-neuronal co-expression of G4C2×36 and GR36 with *USP30* RNAi did not improve the larval locomotor deficit either (Fig. S2F). These data suggest that mitophagy is also not a primary cause but a downstream consequence in *C9orf72* pathogenesis.

### Overexpression of mitochondrial *Sod2* and *catalase* ameliorates *C9orf72* motor phenotypes

Previous studies have shown that targeting oxidative stress may be beneficial in *C9orf72*-related pathology (20). Cellular defence mechanisms such as antioxidants are targeted to different cellular and subcellular locations, due to many sources of ROS. This compartmentalisation also highlights the need for fine tuning of ROS signalling for redox homeostasis as well as the possibility for ROS to signal between compartments (45). Since an increase in mitochondrial ROS was observed in G4C2×36, GR36 and GR1000 flies, we hypothesised that overexpression of antioxidants would suppress behavioural locomotor phenotypes. First, we used a pan-neuronal driver to co-express the major antioxidant genes – cytosolic *Sod1*, mitochondrial *Sod2*, *catalase* (*Cat*) as well as a mitochondrially targeted catalase (*mitoCat*) – with G4C2×36 and GR36. Contrary to our prediction, overexpression of cytosolic *Sod1* significantly worsened the morphology of G4C2×36 and GR36 larvae, becoming even thinner and more developmentally delayed (Fig. 3A). This was reflected by a worsening of the locomotor deficit for G4C2×36 and GR36 larvae, where GR36 larvae co-expressing *Sod1* did not crawl during the assay conditions (Fig. 3B). In contrast, overexpression of mitochondrial *Sod2*, *Cat* or *mitoCat* significantly rescued G4C2×36 and GR36 larval crawling (Fig. 3B). These manipulations also improved larval morphology, particularly for GR36 (Fig. 3A). Exploring the basis for these differential effects we found that expression of *Sod1* was reduced at both mRNA (Fig. S2G) and protein levels (Fig. S2H) in G4C2×36 flies, while *Cat* expression was increased but *Sod2* was unaltered (Fig. S2G).

**Figure 3.**
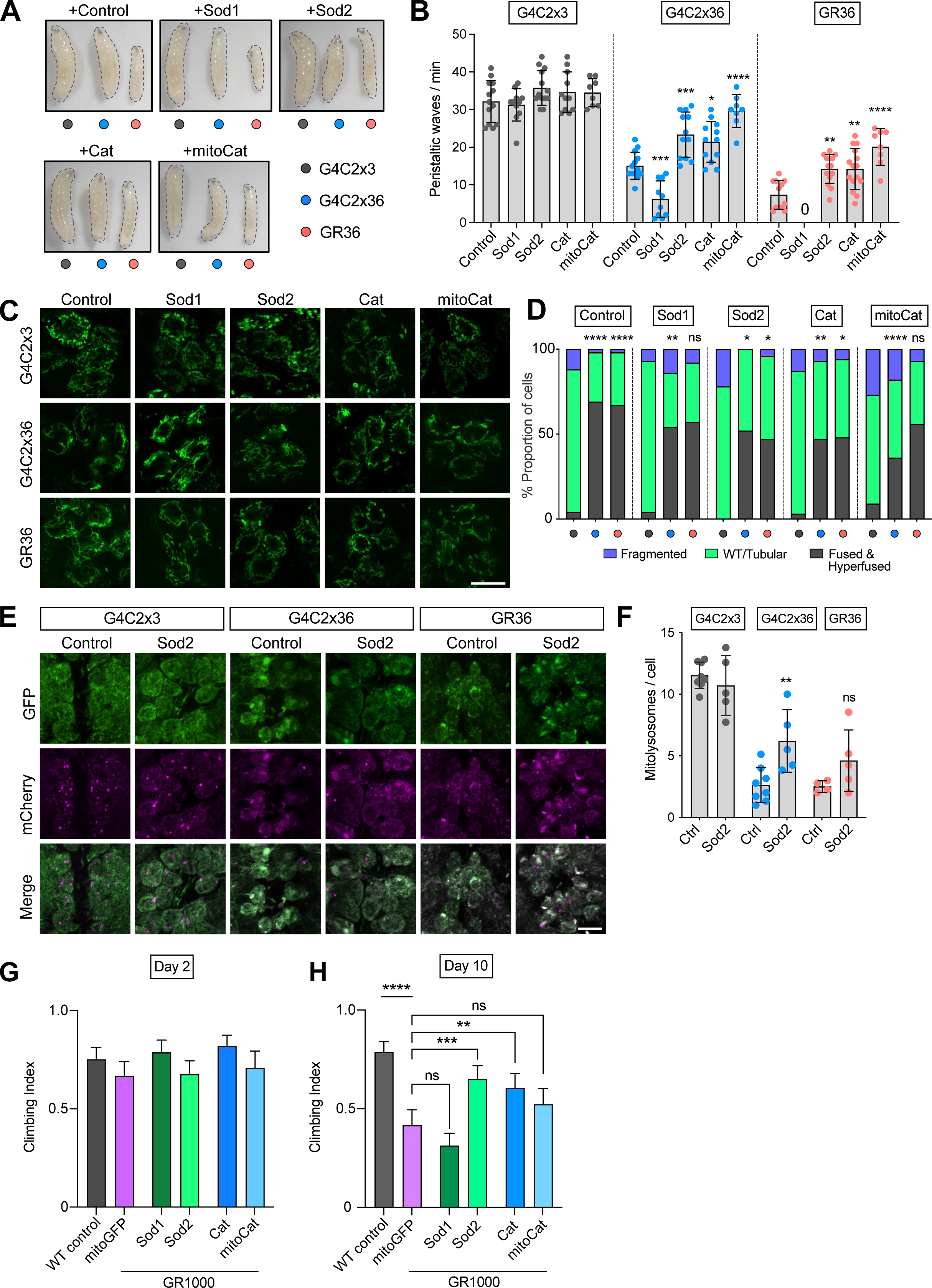
Overexpression of mitochondrial *Sod2* and *catalase* partially rescues *C9orf72* phenotypes. **(A)** Morphology and **(B)** crawling ability of larvae co-expressing G4C2×3, G4C2×36 or GR36 with *LacZ* (control), *Sod1*, *Sod2*, *catalase* (*Cat*) or mitochondrially targeted *catalase* (*mitoCat*) with pan-neuronal driver *nSyb*-GAL4 (mean ± SD; one-way ANOVA with Bonferroni’s multiple comparison test; *p < 0.05, **p < 0.01, ***p < 0.001, ****p < 0.0001; 0 denotes larvae that did not crawl; n = 10-15 larvae). **(C)** Confocal microscopy of larval neurons. Mitochondria are labelled with pan-neuronal expression of mito.GFP, co-expressing G4C2×3, G4C2×36 or GR36 with *nSyb*-GAL4. Genetic manipulations boosting antioxidant capacities by overexpressing *Sod1*, *Sod2*, *Cat* or *mitoCat*. Control is overexpressing *LacZ*. Scale bar = 10 µm. **(D)** Quantification of **C** based on the established scoring system (Chi-squared test; ns = non-significant, *p < 0.05, **p < 0.01, ****p < 0.0001; n = 8-10 larvae). Comparisons for the first *LacZ* group are against G4C2×3, *LacZ*. Otherwise, all the comparisons are against their respective control (*LacZ*) condition. **(E)** Confocal microscopy and **(F)** quantification of the mito-QC mitophagy reporter of larvae co-expressing G4C2×3, G4C2×36 or GR36 with Luciferase RNAi (control) and *Sod2* in the larval ventral ganglion with *nSyb*-GAL4. Scale bar = 10 µm. Mitolysosomes are evident as GFP-negative, mCherry-positive puncta (mean ± SD; one-way ANOVA with Bonferroni’s multiple comparison test; ns = non-significant, **p < 0.01; n = 4-8 larvae). **(G, H)** Climbing analysis of 2- and 10-day-old adults expressing GR1000 co-expressing *mito.GFP*, *Sod1*, *Sod2*, *Cat* and *mitoCat* with *nSyb*-GAL4. WT control is *nSyb/+* (mean ± 95% CI; Kruskal-Wallis non-parametric test with Dunn’s correction; ns = non-significant, **p < 0.01, ***p < 0.001, ****p < 0.0001; n = 60-100 flies).

We next asked whether these manipulations similarly affected the other mitochondrial phenotypes. We found that overexpressing *Sod2*, *Cat* or *mitoCat* significantly reduced the elongated mitochondrial phenotype in G4C2×36 and GR36 larval neurons (Fig. 3C, D). Interestingly, here *Sod1* overexpression also partially rescued this phenotype. Notably, *Sod2* overexpression was also able to partially rescue the defective mitophagy in G4C2×36 larvae, though this did not reach significance for GR36 (Fig. 3E, F). To validate this genetic rescue, we used the GR1000 lines where we also found that *Sod2* or *Cat* overexpression significantly rescued the age-related locomotion defect (Fig. 3G, H), while *mitoCat* expression did not reach significance.

Taken together, amongst all the different genetic manipulations used to reverse mitochondrial phenotypes observed, only overexpression of certain antioxidant enzymes such as mitochondrial *Sod2* and *catalase* were beneficial in rescuing the locomotor deficits. This suggests that oxidative stress is an important upstream pathway in pathogenesis.

### The Keap1/Nrf2 pathway is partially activated in *C9orf72* flies

ROS act as important signalling molecules to coordinate cellular homeostasis and maintain redox balance. Potentially harmful levels of ROS activate redox sensor pathways such as Keap1/Nrf2 which can then upregulate a range of antioxidant genes upon stimulation. Under basal conditions, levels of Nrf2 are kept low by constitutive proteasomal degradation guided by Keap1 functioning as a substate adapter promoting Nrf2 ubiquitination. Upon oxidative stress, the modification of critical reactive cysteine residues on Keap1 leads to released binding of Nrf2, allowing Nrf2 to translocate to the nucleus and activate the expression of a series of antioxidative and cytoprotective genes (23, 24).

*Drosophila* encode homologues of *Keap1* and *Nrf2* (called *cap ‘n’ collar* C isoform, CncC) (31), so we sought to explore whether this pathway is involved in *C9orf72* pathology. First, by assessing the relative nuclear abundance of CncC in neurons of the larval ventral ganglion, we observed an increase in nuclear CncC in both G4C2×36 and GR36 compared to the G4C2×3 control (Fig. 4A, C). We also observed a similar increase in CncC nuclear staining in 5-day-old adult brains of G4C2×36 animals (Fig. 4B, C). This indicates that there is an activation of the Keap1/CncC signalling pathway at early stages which remains activated during disease pathogenesis.

**Figure 4.**
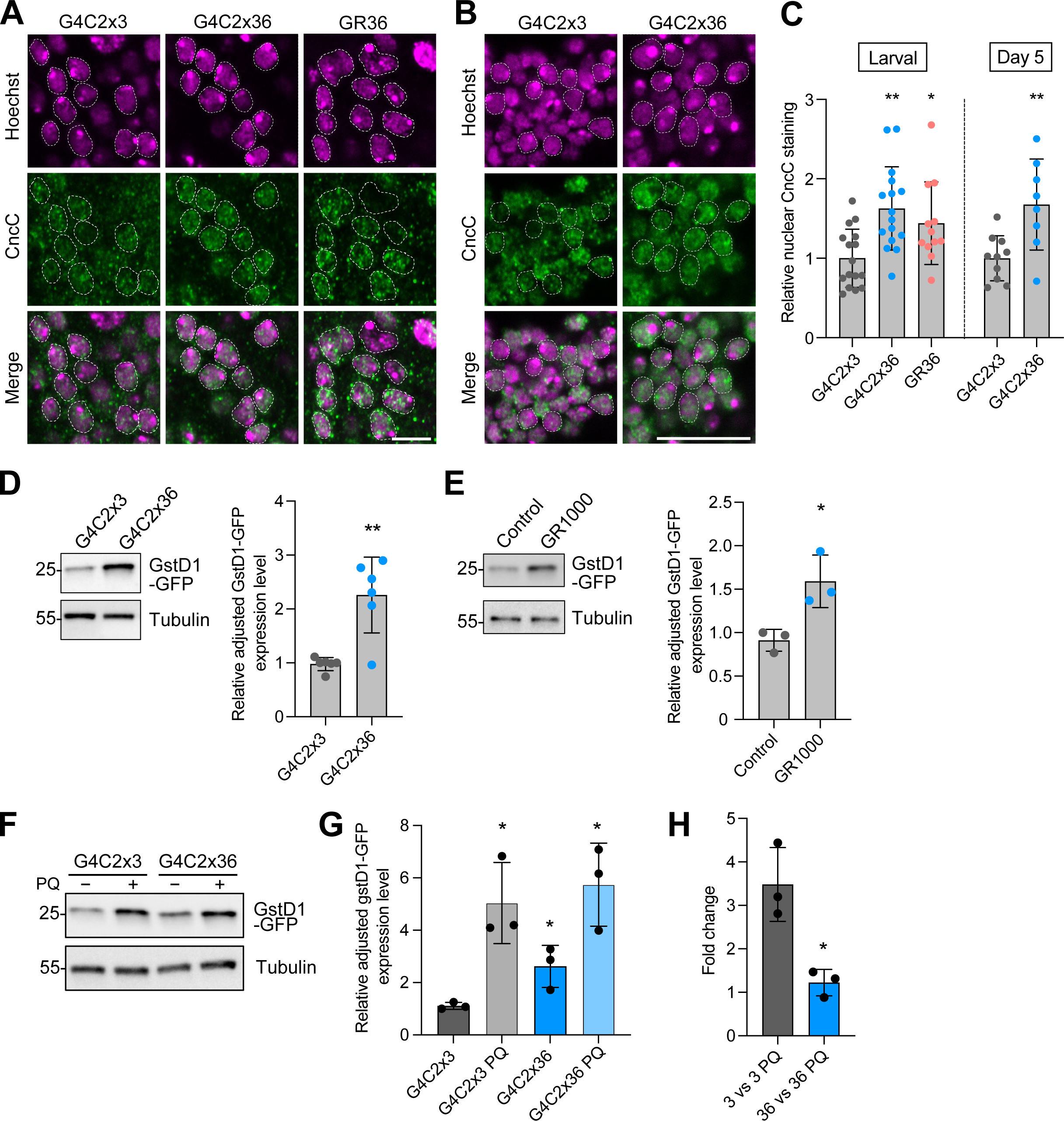
Keap1/CncC pathway is blunted in *C9orf72* pathogenesis. **(A)** Confocal microscopy images of larval ventral nerve ganglion and **(B)** 5-day-old adult brains immunostained for CncC expression, with Hoechst used to identify the nuclei. Pan-neuronal expression of G4C2×36 and GR36 was via *nSyb-*GAL4 and immunostaining intensity was compared to the control, G4C2×3. **(C)** Quantification of relative nuclear staining (mean ± SD; one-way ANOVA with Bonferroni’s multiple comparison test for larval brain analysis; *p < 0.05, ** p < 0.01, n = 12-18 larval brains; unpaired t-test with Welch’s correction for adult brain analysis, ** p < 0.01, n = 8-10 adult brains). **(D, E)** Immunoblotting of pan-neuronal expression using *nSyb*-GAL4 of **(D)** G4C2×3 and G4C2×36 at day 5, and **(E)** LacZ (Control) and GR1000 at day 10, and quantification of *GstD1-GFP* levels, where tubulin was used as the loading control (mean ± SD; unpaired t-test with Welch’s correction; *p < 0.05, ** p < 0.01, n = 3-6 biological replicates). **(F, G)** Immunoblotting of pan-neuronal expression of G4C2×3 and G4C2×36 using *nSyb*-GAL4, in combination with the *GstD1-GFP* reporter, with and without 5 mM PQ treatment. Anti-GFP was used to measure reporter expression at day 5 which was quantified (mean ± SD; one-way ANOVA with Bonferroni’s correction; *p < 0.05, n = 3 biological replicates). **(H)** Chart depicting the fold change of *GstD1-GFP* expression levels comparing G4C2×3 vs G4C2×3 PQ and G4C2×36 vs G4C2×36 PQ (mean ± SD; unpaired t-test with Welch’s correction, * p < 0.05, n = 3 biological replicates).

To further analyse the activity of this signalling pathway, we made use of a reporter transgene which expresses GFP under the control of *GstD1*, an Nrf2 target and prototypical oxidative stress response gene (31), to monitor antioxidant responses. Pan-neuronal expression of both G4C2×36 (Fig. 4D) and GR1000 (Fig. 4E) caused an increase in *GstD1-GFP* levels compared to their respective controls, consistent with an increase in CncC activity. At the same time, we assessed whether G4C2×36 flies could also respond appropriately to ROS-inducing paraquat (PQ) treatment. Again, we saw that the *GstD1-GFP* reporter expression was significantly increased in G4C2×36 flies under basal conditions (− PQ) compared to controls (Fig. 4F, G). However, while *GstD1-GFP* levels further increased upon PQ treatment in both control and G4C2×36 conditions (Fig. 4F, G), this response was proportionally less than the response in a control background (Fig. 4H). These observations, together with the preceding genetic studies, suggest that although the *C9orf72* flies can detect the elevated oxidative stress conditions and mount a response, it appears to be insufficient to confer full protection.

### Genetic and pharmacological inhibition of Keap1 partially rescues *C9orf72* phenotypes

Although the Keap1/Nrf2 pathway was activated in *C9orf72* models, the response seemed to be insufficient. Therefore, we hypothesised that genetically reducing Keap1 levels to constitutively boost CncC activity could benefit *C9orf72* phenotypes as has been seen in a *Drosophila* model of Alzheimer’s disease (46). Indeed, combining a heterozygous mutant of *Keap1* with pan-neuronally driven G4C2×36 and GR36 significantly improved the larval motor ability of both G4C2×36 and GR36 compared to control (Fig. 5A), and visibly improved the larval morphology (Fig. 5B). Furthermore, *Keap1* heterozygosity was also able to fully rescue the GR1000 adult climbing deficit (Fig. 5C).

**Figure 5.**
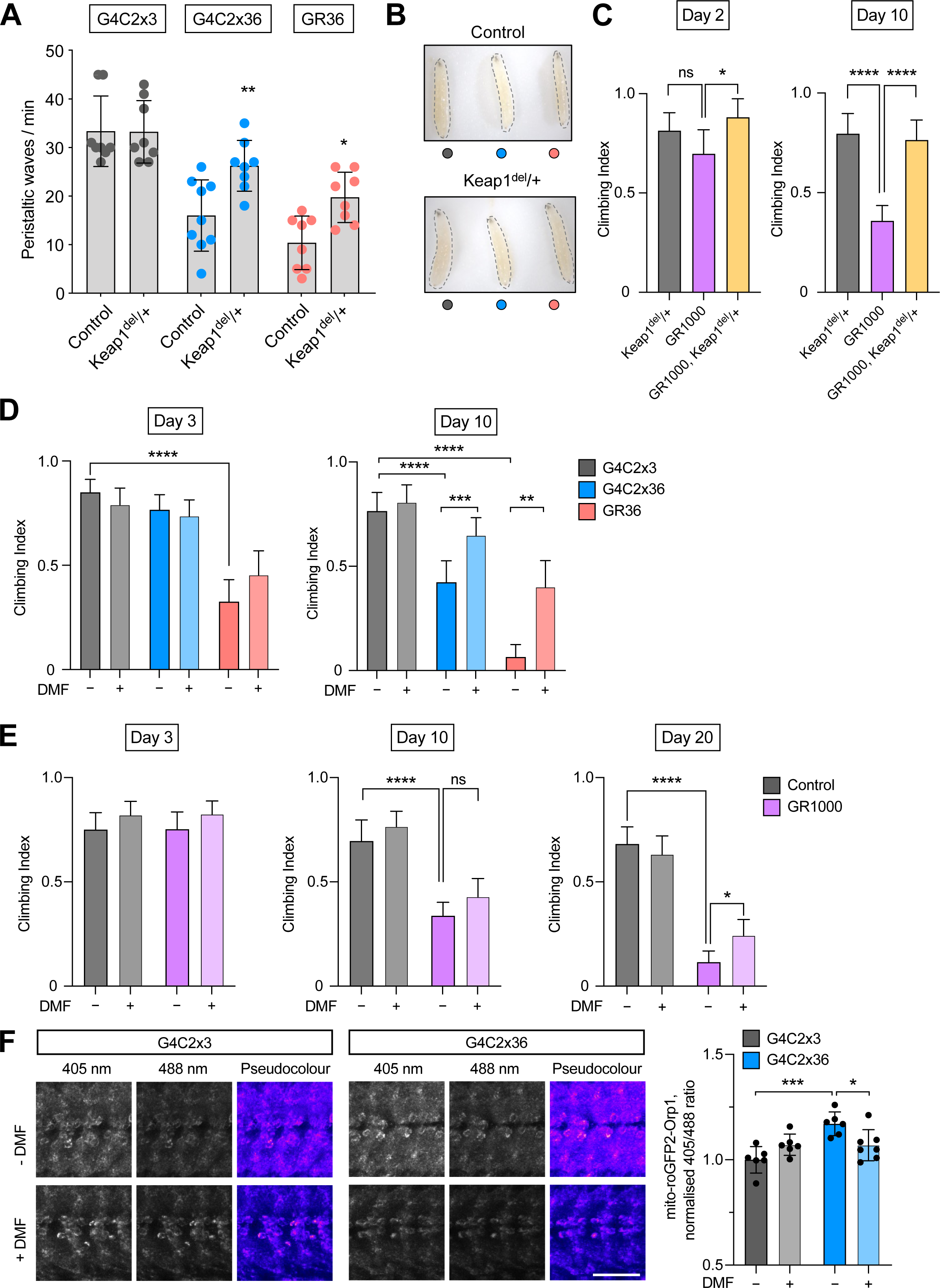
Genetic activation of the Keap1/CncC pathway rescues *C9orf72* toxicity *in vivo*. **(A)** Crawling ability and **(B)** morphology of pan-neuronal expressing G4C2×3, G4C2×36 or GR36 larvae using *nSyb*-GAL4, with heterozygous loss of Keap1 (Keap1^del^/+) compared to a control (WT) background (mean ± SD; one-way ANOVA with Bonferroni’s multiple comparison test; *p < 0.05, **p < 0.01; n = 8-10 larvae). **(C)** Climbing analysis of 2- and 10-day-old adults pan-neuronally (*nSyb*-GAL4) expressing GR1000 with heterozygous loss of Keap1 (mean ± 95% CI; Kruskal-Wallis non-parametric test with Dunn’s correction; *p < 0.05, ****p < 0.0001, ns = non-significant; n = 60-100 flies). **(D)** Climbing analysis of 3- and 10-day-old flies expressing G4C2×3, G4C2×36 and GR36 flies via predominantly motor neuron driver *DIPγ-* GAL4. Flies were raised on food supplemented with either 7 μM dimethyl fumarate (DMF) or ethanol as control (mean ± 95% CI; Kruskal-Wallis non-parametric test with Dunn’s correction; **p < 0.01, ***p < 0.001, ****p < 0.0001; n = 60-100 flies). **(E)** Climbing analysis of 3-, 10- and 20-day-old flies expressing mito.GFP (control) or GR1000 flies via pan-neuronal expression with *nSyb-*GAL4. Flies were raised on food supplemented with either 7 μM DMF or ethanol as control (mean ± 95% CI; Kruskal-Wallis non-parametric test with Dunn’s correction; * p < 0.05, ****p < 0.0001, ns = non-significant; n = 60-100 flies). **(F)** Confocal microscopy of the mito-roGFP2-Orp1 mitochondrial H_2_O_2_ reporter co-expressed with G4C2×3 and G4C2×36 in the larval ventral ganglion with *nSyb*-GAL4. Larvae were subjected to 1 µM DMF or equivalent ethanol control during development. Representative pseudocolour ratio images are depicted and the 405/488 ratio quantified. Scale bar = 50 µm (mean ± SD; one-way ANOVA with Bonferroni’s multiple comparison test; *p < 0.05, *** p < 0.001; n = 6-7 larvae).

The beneficial effects of partial genetic loss of *Keap1* in the *C9orf72* models supports the potential for pharmaceutical agents that modulate Keap1 to also be beneficial in this context. We tested this hypothesis using dimethyl fumarate (DMF), a Keap1-modifying, Nrf2-activating drug with antioxidative and anti-inflammatory properties. DMF acts by the inactivation of Keap1 via succination of its cysteine residues (47), previously shown to exhibit neuroprotective effects in animal models of neurodegeneration (48). In order to assess the effects of extended DMF treatments on motor phenotypes, we expressed G4C2×36 and GR36 via *DIPγ-GAL4* and treated the adult flies with food supplemented with DMF or vehicle (ethanol). After 3 days, G4C2×36 flies did not show a phenotype while the stronger phenotype of GR36 was not significantly improved (Fig. 5D). However, after 10 days of treatment, the decline in locomotor performance in vehicle treated G4C2×36 and GR36 flies was prevented by DMF (Fig. 5D). Similarly, while no benefit was observed at day 3 or day 10 in GR1000 flies with DMF treatment, by day 20, GR1000 flies showed a partial rescue in climbing compared to vehicle control (Fig. 5E).

To test whether DMF treatment improves mitochondrial function in the G4C2×36 model, we employed the mito-roGFP2-Orp1 reporter to measure mitochondrial H_2_O_2_ levels. Here, animals were raised on food containing DMF, or vehicle (ethanol). We again observed an increase in reported H_2_O_2_ levels in G4C2×36 animals which was partially rescued by DMF treatment (Fig. 5F). Taken together, these data indicate that inhibiting Keap1 by DMF treatment can alleviate mitochondrial dysfunction and attenuate *C9orf72* behavioural phenotypes.

### DMF treatment is beneficial in *C9orf72*-ALS/FTD patient-derived neurons

Since we had observed elevated ROS in *C9orf72 Drosophila* models, and found that DMF treatment provided phenotypic benefit, we wanted to test the translatable potential of DMF treatment on patient-relevant material. Fibroblasts from healthy age-matched controls and ALS patients carrying *C9orf72* mutations (Table 1) were reprogrammed into induced neural progenitor cells (iNPCs), then differentiated into induced neurons (iNeurons) (49). Assessing the level of oxidative stress in these cells using MitoSOX, we found a modest increase in mitochondrial ROS in *C9orf72* iNeurons (Fig. 6A, B), in line with that observed in *Drosophila*. We also found that there was a higher level of nuclear Nrf2 in patient iNeurons (Fig. 6C, D) mirroring the *C9orf72 Drosophila* phenotypes.

**Figure 6.**
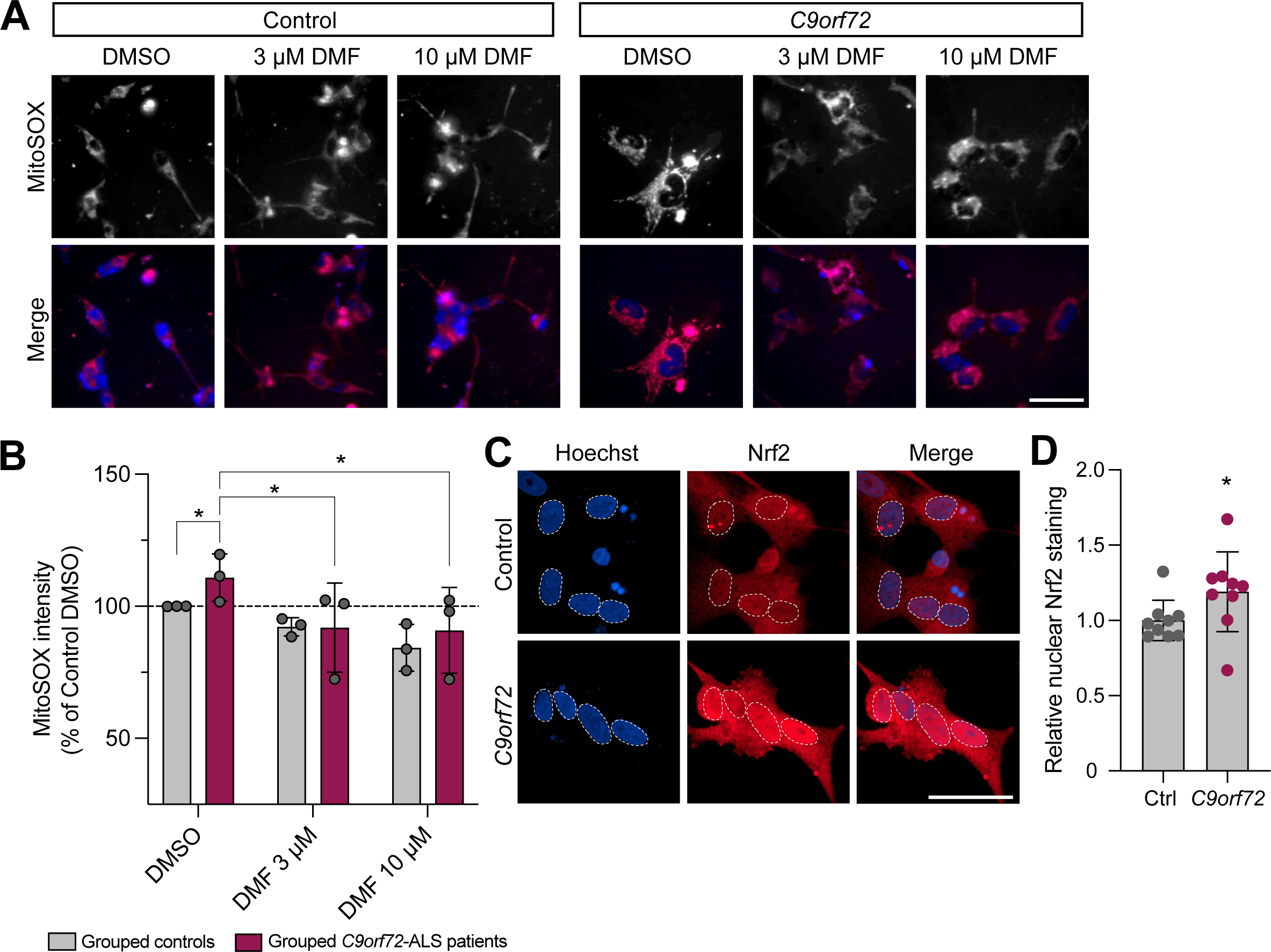
DMF activation of the Keap1/Nrf2 pathway reduces mitochondrial ROS in *C9orf72* patient-derived iNeurons. **(A)** MitoSOX staining of *C9orf72* iNeurons compared with healthy controls treated with 3 μM and 10 μM DMF or DMSO as control. Scale bar = 50 µm. **(B)** Quantification of MitoSOX staining (mean ± SD; two-way ANOVA with Dunnett’s multiple comparison test; *p < 0.05, n = 3 age-matched patients vs control). **(C)** Confocal microscopy images of *C9orf72* iNeurons compared with healthy controls immunostained for Nrf2 expression, with Hoechst used to identify the nuclei. **(D)** Quantification of relative nuclear staining (mean ± SD; unpaired t-test with Welch’s correction; *p < 0.05, n = 3 age-matched patients vs control).

Testing two different concentrations, 24-hour treatment with 3 μM or 10 μM DMF was able to reduce the mitochondrial ROS levels in *C9orf72* iNeurons (Fig. 6A, B). In comparison, we analysed as a positive control treatment with edaravone (EDV), a known ROS scavenger and FDA-approved ALS drug, and observed that 24-hour treatment with 30 μM and 100 μM EDV reduced ROS levels in *C9orf72* iNeurons to a similar extent as DMF (Fig. S3A, B). Although significant, the effect size of DMF or EDV treatment is small and the impact on neuronal function or survival is currently unknown. Therefore, more investigation is required to validate these initial results. Nevertheless, taken together, these data suggest that activating the Keap1/Nrf2 pathway by using therapeutic interventions such as DMF, can be beneficial across multiple model systems including *in vivo C9orf72 Drosophila* models and human *C9orf72* patient-derived neurons.

## Discussion

We have conducted a comprehensive characterisation of mitochondrial dysfunction using three different models of *C9orf72* ALS/FTD and found disrupted morphology with hyperfused mitochondria, reduced mitophagy, impaired respiration and increased mitochondrial ROS production, all in a neuronal context *in vivo*. Genetic interaction studies showed that only overexpression of mitochondrial *Sod2* and *catalase* were able to significantly rescue *C9orf72* behavioural phenotypes, and further rescued other mitochondrial deficits such as mitophagy. Together, these data suggest a causative link between mitochondrial dysfunction, ROS and behavioural phenotypes.

ROS has been well-characterised in the context of ALS (38), however, it is important to understand the relative contributions of different ROS species in different cellular compartments in order to identify more targeted treatment options. Addressing this, we have observed a robust increase in mitochondrial ROS whereas the effect on cytosolic ROS was more variable or unaltered. Consistent with the compartmentalised effects, we found that overexpressing mitochondrial *Sod2* or a mitochondrially-targeted *Cat* partially rescued *C9orf72* phenotypes, as did overexpression of cytosolic *Cat*, which suggests that mitochondrial superoxide produced is rapidly dismutated causing a cytosolic effect hence both cytosolic and mitochondrially-targeted *Cat* is beneficial.

We were surprised to find that *Sod1* overexpression exacerbated *C9orf72* phenotypes, contrary to previous findings from Lopez-Gonzalez et al. (20). SOD1 is a cytosolic ubiquitous enzyme with several functions, primarily involved in scavenging superoxide as well as modulating cellular respiration, energy metabolism and posttranslational modifications (50). Mutations in *SOD1* also cause ALS, most likely via misfolding of the protein, causing a toxic gain of function, although the exact mechanism leading to motor neuron death is still elusive (51). Although we observe reduced levels of *Sod1* mRNA transcripts and protein, silencing *SOD1* in mice does not itself cause neurodegeneration (52). However, it would be interesting to investigate whether Sod1 misfolding or aggregation may contribute to *C9orf72* pathogenesis. Indeed, Forsberg et al. (53) found abundant inclusions containing misfolded WT SOD1 were found in spinal and cortical motor neurons from patients carrying mutations in *C9orf72*. Taken together, these findings highlight the importance of studying compartmentalised ROS effects in the context of disease pathology.

While defective autophagy has previously been noted in *C9orf72 Drosophila* and other models (54), we extend these observations to provide novel *in vivo* evidence that mitophagy is also perturbed. However, a mechanistic link between disrupted mitophagy and the upstream oxidative stress and modulation of the Nrf2 pathway is lacking. Nrf2 activity is known to promote mitochondrial homeostasis at multiple levels including biogenesis and turnover (55). Notably, the autophagy adapter p62 both competes with Nrf2 for Keap1 binding (56) and is an Nrf2 target gene, thereby creating a positive feedback loop (56, 57). Moreover, Nrf2 activation was shown to induce mitophagy independently of the Pink1/parkin pathway, and rescue *Pink1/parkin* phenotypes *in vivo* (58, 59). Therefore, it will be interesting to investigate a potential link between *C9orf72*–Nrf2/CncC–p62/ref(2)P–mitophagy/autophagy as it may provide mechanistic insight, in all our model systems.

The protective role of the well-known antioxidant and cytoprotective Keap1/Nrf2 pathway has been discussed as a therapeutic target for treatment against many neurodegenerative diseases including ALS (24). Investigating this, we found increased nuclear localisation of Nrf2 and concomitant upregulation of the *GstD1-GFP* reporter, a proxy for Nrf2 activation, indicating that the pathway is upregulated upon *C9orf72* toxicity. However, despite the initiated activation, this was clearly insufficient to prevent excessive ROS and the resulting neurotoxicity which could be prevented by transgenic overexpression of antioxidant enzymes. While the reasons behind the blunted pathway activation are unclear, it is encouraging that both genetic reduction of Keap1, as well as its pharmacological targeting with DMF, could suppress *C9orf72* toxicity *in vivo*, and in patient-derived iNeurons.

While some studies have reported the Nrf2 pathway to be disrupted in ALS patient samples (60–62), very limited data exists on *C9orf72*-related ALS samples. Recently, Jimenez-Villegas et al. (63) have shown that Nrf2 activation is impaired in cell culture models of arginine-DPR toxicity, where they also saw an improvement in cell viability upon treatment with DMF. We have extended these findings showing the benefit of targeting Keap1/Nrf2 *in vivo* as well as patient-derived iNeurons which further emphasises the importance of Nrf2 activation as a potential therapeutic target. Encouragingly, a recent phase 2 clinical trial to test the efficacy of DMF in sporadic ALS patients was conducted (64). However, while the study concluded there was efficacy on the primary endpoint (Amyotrophic Lateral Sclerosis Functional Rating Scale-Revised (ALSFRS-R) score), there was a reduced decline in neurophysiological index, suggesting preservation of lower motor neuron function. The authors also noted that participants showed an ‘unusually slow disease progression’ and that a larger trial is needed for verification (64).

In conclusion, our results provide compelling evidence that mitochondrial oxidative stress is an important upstream pathogenic mechanism leading to downstream mitochondrial dysfunction such as alterations in mitochondrial function and turnover. Consequently, targeting one of the main intracellular defence mechanisms to counteract oxidative stress – the Keap1/Nrf2 signalling pathway – could be a viable therapeutic strategy for ALS/FTD. While DMF treatment shows promise, more research is needed to understand the underlying mechanisms behind disease pathogenesis and progression.

## List of Abbreviations

ALS: Amyotrophic Lateral Sclerosis
ALSFRS-R: Amyotrophic Lateral Sclerosis Functional Rating Scale-Revised
C9orf72: Chromosome 9 Open Reading Frame 72
Cat: Catalase
DMF: Dimethyl fumarate
DPR: Dipeptide repeat proteins
EDV: Edaravone
FTD: Frontotemporal Dementia
GR: Glycine-arginine
Grx1: Glutaredoxin 1
GSSG: Oxidised glutathione
H2O2: Hydrogen peroxide
Nrf2: Nuclear factor erythroid 2-related factor 2
OMM: Outer mitochondrial membrane
RAN: Repeat-associated non-AUG translation
roGFP2: Redox-sensitive fluorescent protein
ROS: Reactive Oxygen Species

## Declarations

### Ethics approval and consent to participate

For the iNeuron work, informed consent was obtained from all participants before collection of fibroblast biopsies (Sheffield Teaching Hospital (STH) study number: STH16573, Research Ethics Committee reference: 12/YH/0330); fibroblasts obtained by Cedars-Sinai are covered by MTA. *Drosophila* stocks were obtained from the Bloomington *Drosophila* Stock Center which is supported by grant NIH P40OD018537. Experimental work with Drosophila is exempt from ethical approval as they are not covered by the Animals (Scientific Procedures) Act 1986.

### Consent for publication

Not applicable

### Availability of data and materials

All data needed to evaluate the conclusions in the paper are present in the paper and/or the Supplementary Materials. This study includes no data deposited in external repositories. Additional data related to this paper may be requested from the authors.

### Competing interests

The authors declare that they have no competing interests.

### Funding

This work was supported by Medical Research Council (MRC) core funding (MC_UU_00015/6 and MC_UU_0028/6) to AJW, MRC studentships (MC_ST_U18009) to WHA and MJT, and MRC project grants; MR/V003933/1 to AJW/LMF and MR/W00416X/1 to LF. JAKL was supported by a Battelle Memorial Institute Wadsworth PhD Fellowship. HM/JAKL are supported by MNDA 943-793. HM was supported by Parkinson’s UK Senior Research Fellowship F-1301. This is independent research funded by the above funders and partially carried out at the National Institute for Health and Care Research (NIHR) Sheffield Biomedical Research Centre (BRC). The views expressed are those of the author(s) and not necessarily those of the funders, the NIHR or the Department of Health and Social Care.

## Authors’ contributions

WHA: Conceptualisation, Methodology, Formal analysis, Investigation, Writing - Original Draft,

Writing - Review & Editing, Visualisation

LMF: Methodology, Investigation, Writing - Review & Editing

ASM: Methodology, Supervision, Writing - Review & Editing

JAKL: Methodology, Formal analysis, Investigation, Writing - Review & Editing,

MJT: Methodology, Investigation, Writing - Review & Editing,

HAP: Conceptualisation, Methodology, Writing - Review & Editing

SG: Methodology

KR: Methodology

LF: Methodology, Supervision, Writing - Review & Editing

HM: Conceptualisation, Formal analysis, Writing – Review & Editing, Supervision, Funding acquisition

AJW: Conceptualisation, Validation, Formal analysis, Writing - Original Draft, Writing – Review & Editing, Supervision, Project administration, Funding acquisition

## Acknowledgements

We kindly thank the following for generously sharing fly lines: Dr Robert Carrillo, Prof. Adrian Isaacs, Dr Ryan West, Prof. Alberto Sanz, Prof. Linda Partridge as well as Dr Fengwei Yu for sharing the CncC antibody. We also extend our thanks to Dr Scott Allen for additional supervision of JAKL and joint acquisition of the PhD Fellowship. Finally, we thank all members of the Whitworth lab for discussions and feedback on the manuscript.

## Figure Legends

**Supplementary Figure 1. *C9orf72 Drosophila* models show impaired mitochondrial function and autophagy but have normal cytosolic ROS levels**

**(A)** Lifespan curves of G4C2×36 and GR36 flies compared with G4C2×3 control driven by *Dipγ-*GAL4. Mantel-Cox log-rank test (n ≥ 80 flies). **(B,C)** Survival analysis or flies exposed to **(B)** 10 mM paraquat (PQ) and **(C)** 1% H_2_O_2_ treatment. Mantel-Cox log-rank test (n ≥ 60 flies). **(C)** Confocal microscopy of the cyto-Grx1-roGFP2 cytosolic glutathione redox potential reporter co-expressed with G4C2×3, G4C2×36 and GR36 in the larval ventral ganglion with *nSyb*-GAL4. Representative pseudocolour ratio images are depicted and the 405/488 ratio quantified. Scale bar = 50 µm (mean ± SD; one-way ANOVA with Bonferroni’s multiple comparison test; ns = non-significant n = 9-10 larvae). **(E)** Confocal microscopy analysis of DHE staining in G4C2×3, G4C2×36 and GR36 larval ventral ganglion using *nSyb*-GAL4 where DHE intensity is quantified and normalised to the mean of the G4C2×3 control (mean ± SD; one-way ANOVA with Bonferroni’s multiple comparison test; ns = non-significant n = 8-13 larvae). **(F)** Confocal microscopy of the GFP-mCherry-Atg8a autophagy flux reporter pan-neuronally co-expressing G4C2×3, G4C2×36 and GR36 in the larval ventral ganglion using *nSyb*-GAL4. Scale bar = 10 µm. Autolysosomes are quantified as mCherry-positive puncta (mean ± SD; one-way ANOVA with Bonferroni’s multiple comparison test; *p < 0.05, **p < 0.01; n = 8-9 larvae). **(G)** Western blot analysis of 5-day-old pan-neuronally expressing G4C2×3 and G4C2×36 fly brains using *nSyb*-GAL4. Immunoblots were probed with GABARAP to measure Atg8a levels and ref(2)P where tubulin was used as a loading control. A low and high exposure for Atg8a is depicted, where high exposure shows the lipidated Atg8a-II band clearer. **(H)** Quantification of ref(2)P levels (mean ± SD; unpaired t-test with Welch’s corrections; **p < 0.01; n = 5 biological replicates).

**Supplementary Figure 2. Genetic manipulation of mitochondrial dynamics or mitophagy does not rescue *C9orf72* phenotypes**

**(A)** Confocal microscopy of larval neurons where mitochondria are labelled with pan-neuronal expression of mito.GFP, co-expressing G4C2×3, G4C2×36 or GR36 using *nSyb*-GAL4. Genetic manipulations promoting fission by overexpressing *LacZ* (control), *Tango11* and *Drp1* as well as reducing fusion by using heterozygous loss-of-function mutations of *Opa1* and *Marf*. Scale bar = 10 µm. (**B)** Quantification of **A** based on the established scoring system (Chi-squared test; ns = non-significant, *p < 0.05, **p < 0.01, ***p < 0.001, ****p < 0.0001; n = 6-12. Comparisons for the first *LacZ* group are against G4C2×3, *LacZ*. Otherwise, all the comparisons are against their respective control (*LacZ*) condition). **(C)** Larval crawling of G4C2×3, G4C2×36 and GR36 larvae pan-neuronally co-expressing pro fission factors *Tango11* and *Drp1* as well as heterozygous reduction of pro fusion factors *Opa1* and *Marf* (mean ± SD; one-way ANOVA with Bonferroni’s multiple comparison test; *p < 0.05, **p < 0.01; n = 6-12 larvae). **(D, E)** Confocal microscopy of the mito-QC mitophagy reporter of G4C2×3, G4C2×36 and GR36 larvae co-expressing Luciferase RNAi (control) and *USP30* RNAi with *nSyb*-GAL4. Scale bar = 10 µm. (mean ± SD; one-way ANOVA with Bonferroni’s multiple comparison test; ns = non-significant, ***p < 0.001; n = 4-8 larvae). **(F)** Larval crawling of G4C2×3, G4C2×36 and GR36 larvae co-expressing Luciferase RNAi (control) and USP30 RNAi with *nSyb*-GAL4 (mean ± SD; one-way ANOVA with Bonferroni’s multiple comparison test; data not significant, n = 6-12 larvae). **(G)** qRT-PCR was performed using 5-day-old adult heads pan-neuronally expressing G4C2×3 and G4C2×36 with *nSyb*-GAL4, analysing basal mRNA transcript levels of *Sod1*, *Sod2* and *catalase* (*Cat*) (mean ± SD; unpaired t-test with Welch’s correction; *p < 0.05, n = 4 biological replicates). **(H)** Quantification of Sod1 protein levels (mean ± SD; unpaired t-test with Welch’s corrections; *p < 0.05, n = 4 biological replicates).

**Supplementary Figure 3. Edaravone treatment reduces mitochondrial ROS in *C9orf72* patient-derived iNeurons**

**(A)** MitoSOX staining of *C9orf72* iNeurons compared with healthy controls treated with 30 μM and 100 μM of edaravone (EDV) or DMSO as control. Scale bar = 50 µm. **(B)** Quantification of MitoSOX staining (mean ± SD; two-way ANOVA with Dunnett’s multiple comparison test; *p < 0.05, **p < 0.01, n = 3 age-matched patients vs control).

## References

1. Burrell JR, Halliday GM, Kril JJ, Ittner LM, Gotz J, Kiernan MC, et al. The frontotemporal dementia-motor neuron disease continuum. Lancet. 2016;388(10047):919–31.

2. Masrori P, Van Damme P. Amyotrophic lateral sclerosis: a clinical review. Eur J Neurol. 2020;27(10):1918–29.

3. DeJesus-Hernandez M, Mackenzie IR, Boeve BF, Boxer AL, Baker M, Rutherford NJ, et al. Expanded GGGGCC hexanucleotide repeat in noncoding region of C9ORF72 causes chromosome 9p-linked FTD and ALS. Neuron. 2011;72(2):245–56.

4. Renton AE, Majounie E, Waite A, Simon-Sanchez J, Rollinson S, Gibbs JR, et al. A hexanucleotide repeat expansion in C9ORF72 is the cause of chromosome 9p21-linked ALS-FTD. Neuron. 2011;72(2):257–68.

5. Balendra R, Isaacs AM. C9orf72-mediated ALS and FTD: multiple pathways to disease. Nat Rev Neurol. 2018;14(9):544–58.

6. Mizielinska S, Gronke S, Niccoli T, Ridler CE, Clayton EL, Devoy A, et al. C9orf72 repeat expansions cause neurodegeneration in Drosophila through arginine-rich proteins. Science. 2014;345(6201):1192–4.

7. Kwon I, Xiang S, Kato M, Wu L, Theodoropoulos P, Wang T, et al. Poly-dipeptides encoded by the C9orf72 repeats bind nucleoli, impede RNA biogenesis, and kill cells. Science. 2014;345(6201):1139–45.

8. Wen X, Tan W, Westergard T, Krishnamurthy K, Markandaiah SS, Shi Y, et al. Antisense proline-arginine RAN dipeptides linked to C9ORF72-ALS/FTD form toxic nuclear aggregates that initiate in vitro and in vivo neuronal death. Neuron. 2014;84(6):1213–25.

9. Freibaum BD, Lu Y, Lopez-Gonzalez R, Kim NC, Almeida S, Lee KH, et al. GGGGCC repeat expansion in C9orf72 compromises nucleocytoplasmic transport. Nature. 2015;525(7567):129–33.

10. Jovicic A, Mertens J, Boeynaems S, Bogaert E, Chai N, Yamada SB, et al. Modifiers of C9orf72 dipeptide repeat toxicity connect nucleocytoplasmic transport defects to FTD/ALS. Nat Neurosci. 2015;18(9):1226–9.

11. Boeynaems S, Bogaert E, Michiels E, Gijselinck I, Sieben A, Jovicic A, et al. Drosophila screen connects nuclear transport genes to DPR pathology in c9ALS/FTD. Sci Rep. 2016;6:20877.

12. Zhang K, Donnelly CJ, Haeusler AR, Grima JC, Machamer JB, Steinwald P, et al. The C9orf72 repeat expansion disrupts nucleocytoplasmic transport. Nature. 2015;525(7567):56–61.

13. Baldwin KR, Godena VK, Hewitt VL, Whitworth AJ. Axonal transport defects are a common phenotype in Drosophila models of ALS. Hum Mol Genet. 2016;25(12):2378–92.

14. Solomon DA, Stepto A, Au WH, Adachi Y, Diaper DC, Hall R, et al. A feedback loop between dipeptide-repeat protein, TDP-43 and karyopherin-alpha mediates C9orf72-related neurodegeneration. Brain. 2018;141(10):2908–24.

15. Nguyen L, Montrasio F, Pattamatta A, Tusi SK, Bardhi O, Meyer KD, et al. Antibody Therapy Targeting RAN Proteins Rescues C9 ALS/FTD Phenotypes in C9orf72 Mouse Model. Neuron. 2020;105(4):645–62 e11.

16. Dafinca R, Barbagallo P, Talbot K. The Role of Mitochondrial Dysfunction and ER Stress in TDP-43 and C9ORF72 ALS. Front Cell Neurosci. 2021;15:653688.

17. Dafinca R, Scaber J, Ababneh N, Lalic T, Weir G, Christian H, et al. C9orf72 Hexanucleotide Expansions Are Associated with Altered Endoplasmic Reticulum Calcium Homeostasis and Stress Granule Formation in Induced Pluripotent Stem Cell-Derived Neurons from Patients with Amyotrophic Lateral Sclerosis and Frontotemporal Dementia. Stem Cells. 2016;34(8):2063–78.

18. Onesto E, Colombrita C, Gumina V, Borghi MO, Dusi S, Doretti A, et al. Gene-specific mitochondria dysfunctions in human TARDBP and C9ORF72 fibroblasts. Acta Neuropathol Commun. 2016;4(1):47.

19. Mehta AR, Gregory JM, Dando O, Carter RN, Burr K, Nanda J, et al. Mitochondrial bioenergetic deficits in C9orf72 amyotrophic lateral sclerosis motor neurons cause dysfunctional axonal homeostasis. Acta Neuropathol. 2021;141(2):257–79.

20. Lopez-Gonzalez R, Lu Y, Gendron TF, Karydas A, Tran H, Yang D, et al. Poly(GR) in C9ORF72-Related ALS/FTD Compromises Mitochondrial Function and Increases Oxidative Stress and DNA Damage in iPSC-Derived Motor Neurons. Neuron. 2016;92(2):383–91.

21. Choi SY, Lopez-Gonzalez R, Krishnan G, Phillips HL, Li AN, Seeley WW, et al. C9ORF72-ALS/FTD-associated poly(GR) binds Atp5a1 and compromises mitochondrial function in vivo. Nat Neurosci. 2019;22(6):851–62.

22. Li S, Wu Z, Tantray I, Li Y, Chen S, Dong J, et al. Quality-control mechanisms targeting translationally stalled and C-terminally extended poly(GR) associated with ALS/FTD. Proc Natl Acad Sci U S A. 2020;117(40):25104–15.

23. Hayes JD, Dinkova-Kostova AT. The Nrf2 regulatory network provides an interface between redox and intermediary metabolism. Trends Biochem Sci. 2014;39(4):199–218.

24. Jimenez-Villegas J, Ferraiuolo L, Mead RJ, Shaw PJ, Cuadrado A, Rojo AI. NRF2 as a therapeutic opportunity to impact in the molecular roadmap of ALS. Free Radic Biol Med. 2021;173:125–41.

25. Carrillo RA, Ozkan E, Menon KP, Nagarkar-Jaiswal S, Lee PT, Jeon M, et al. Control of Synaptic Connectivity by a Network of Drosophila IgSF Cell Surface Proteins. Cell. 2015;163(7):1770–82.

26. Venkatasubramanian L, Guo Z, Xu S, Tan L, Xiao Q, Nagarkar-Jaiswal S, et al. Stereotyped terminal axon branching of leg motor neurons mediated by IgSF proteins DIP-alpha and Dpr10. Elife. 2019;8.

27. West RJH, Sharpe JL, Voelzmann A, Munro AL, Hahn I, Baines RA, et al. Co-expression of C9orf72 related dipeptide-repeats over 1000 repeat units reveals age- and combination-specific phenotypic profiles in Drosophila. Acta Neuropathol Commun. 2020;8(1):158.

28. Radyuk SN, Rebrin I, Klichko VI, Sohal BH, Michalak K, Benes J, et al. Mitochondrial peroxiredoxins are critical for the maintenance of redox state and the survival of adult Drosophila. Free Radic Biol Med. 2010;49(12):1892–902.

29. Park J, Lee G, Chung J. The PINK1-Parkin pathway is involved in the regulation of mitochondrial remodeling process. Biochem Biophys Res Commun. 2009;378(3):518–23.

30. Sandoval H, Yao CK, Chen K, Jaiswal M, Donti T, Lin YQ, et al. Mitochondrial fusion but not fission regulates larval growth and synaptic development through steroid hormone production. Elife. 2014;3.

31. Sykiotis GP, Bohmann D. Keap1/Nrf2 signaling regulates oxidative stress tolerance and lifespan in Drosophila. Dev Cell. 2008;14(1):76–85.

32. Castillo-Quan JI, Li L, Kinghorn KJ, Ivanov DK, Tain LS, Slack C, et al. Lithium Promotes Longevity through GSK3/NRF2-Dependent Hormesis. Cell Rep. 2016;15(3):638–50.

33. Lee JJ, Sanchez-Martinez A, Martinez Zarate A, Beninca C, Mayor U, Clague MJ, et al. Basal mitophagy is widespread in Drosophila but minimally affected by loss of Pink1 or parkin. J Cell Biol. 2018;217(5):1613–22.

34. Stewart BA, Atwood HL, Renger JJ, Wang J, Wu CF. Improved stability of Drosophila larval neuromuscular preparations in haemolymph-like physiological solutions. J Comp Physiol A. 1994;175(2):179–91.

35. Pfaffl MW. A new mathematical model for relative quantification in real-time RT-PCR. Nucleic Acids Res. 2001;29(9):e45.

36. Meyer K, Ferraiuolo L, Miranda CJ, Likhite S, McElroy S, Renusch S, et al. Direct conversion of patient fibroblasts demonstrates non-cell autonomous toxicity of astrocytes to motor neurons in familial and sporadic ALS. Proc Natl Acad Sci U S A. 2014;111(2):829–32.

37. Gatto N, Dos Santos Souza C, Shaw AC, Bell SM, Myszczynska MA, Powers S, et al. Directly converted astrocytes retain the ageing features of the donor fibroblasts and elucidate the astrocytic contribution to human CNS health and disease. Aging Cell. 2021;20(1):e13281.

38. Smith EF, Shaw PJ, De Vos KJ. The role of mitochondria in amyotrophic lateral sclerosis. Neurosci Lett. 2019;710:132933.

39. Brand AH, Perrimon N. Targeted gene expression as a means of altering cell fates and generating dominant phenotypes. Development. 1993;118(2):401–15.

40. Li S, Wu Z, Li Y, Tantray I, De Stefani D, Mattarei A, et al. Altered MICOS Morphology and Mitochondrial Ion Homeostasis Contribute to Poly(GR) Toxicity Associated with C9-ALS/FTD. Cell Rep. 2020;32(5):107989.

41. Albrecht SC, Barata AG, Grosshans J, Teleman AA, Dick TP. In vivo mapping of hydrogen peroxide and oxidized glutathione reveals chemical and regional specificity of redox homeostasis. Cell Metab. 2011;14(6):819–29.

42. Cunningham KM, Maulding K, Ruan K, Senturk M, Grima JC, Sung H, et al. TFEB/Mitf links impaired nuclear import to autophagolysosomal dysfunction in C9-ALS. Elife. 2020;9.

43. Bingol B, Tea JS, Phu L, Reichelt M, Bakalarski CE, Song Q, et al. The mitochondrial deubiquitinase USP30 opposes parkin-mediated mitophagy. Nature. 2014;510(7505):370–5.

44. Sanchez-Martinez A, Martinez A, Whitworth AJ. FBXO7/ntc and USP30 antagonistically set the ubiquitination threshold for basal mitophagy and provide a target for Pink1 phosphorylation in vivo. PLoS Biol. 2023;21(8):e3002244.

45. Wang Y, Branicky R, Noe A, Hekimi S. Superoxide dismutases: Dual roles in controlling ROS damage and regulating ROS signaling. J Cell Biol. 2018;217(6):1915–28.

46. Kerr F, Sofola-Adesakin O, Ivanov DK, Gatliff J, Gomez Perez-Nievas B, Bertrand HC, et al. Direct Keap1-Nrf2 disruption as a potential therapeutic target for Alzheimer’s disease. PLoS Genet. 2017;13(3):e1006593.

47. Yamamoto M, Kensler TW, Motohashi H. The KEAP1-NRF2 System: a Thiol-Based Sensor-Effector Apparatus for Maintaining Redox Homeostasis. Physiol Rev. 2018;98(3):1169–203.

48. Majkutewicz I. Dimethyl fumarate: A review of preclinical efficacy in models of neurodegenerative diseases. Eur J Pharmacol. 2022;926:175025.

49. Castelli LM, Lin YH, Sanchez-Martinez A, Gul A, Mohd Imran K, Higginbottom A, et al. A cell-penetrant peptide blocking C9ORF72-repeat RNA nuclear export reduces the neurotoxic effects of dipeptide repeat proteins. Sci Transl Med. 2023;15(685):eabo3823.

50. Saccon RA, Bunton-Stasyshyn RK, Fisher EM, Fratta P. Is SOD1 loss of function involved in amyotrophic lateral sclerosis? Brain. 2013;136(Pt 8):2342–58.

51. Cenini G, Lloret A, Cascella R. Oxidative Stress in Neurodegenerative Diseases: From a Mitochondrial Point of View. Oxid Med Cell Longev. 2019;2019:2105607.

52. Reaume AG, Elliott JL, Hoffman EK, Kowall NW, Ferrante RJ, Siwek DF, et al. Motor neurons in Cu/Zn superoxide dismutase-deficient mice develop normally but exhibit enhanced cell death after axonal injury. Nat Genet. 1996;13(1):43–7.

53. Forsberg K, Graffmo K, Pakkenberg B, Weber M, Nielsen M, Marklund S, et al. Misfolded SOD1 inclusions in patients with mutations in C9orf72 and other ALS/FTD-associated genes. J Neurol Neurosurg Psychiatry. 2019;90(8):861–9.

54. Diab R, Pilotto F, Saxena S. Autophagy and neurodegeneration: Unraveling the role of C9ORF72 in the regulation of autophagy and its relationship to ALS-FTD pathology. Front Cell Neurosci. 2023;17:1086895.

55. Holmstrom KM, Kostov RV, Dinkova-Kostova AT. The multifaceted role of Nrf2 in mitochondrial function. Curr Opin Toxicol. 2016;1:80–91.

56. Komatsu M, Kurokawa H, Waguri S, Taguchi K, Kobayashi A, Ichimura Y, et al. The selective autophagy substrate p62 activates the stress responsive transcription factor Nrf2 through inactivation of Keap1. Nat Cell Biol. 2010;12(3):213–23.

57. Jain A, Rusten TE, Katheder N, Elvenes J, Bruun JA, Sjottem E, et al. p62/Sequestosome-1, Autophagy-related Gene 8, and Autophagy in Drosophila Are Regulated by Nuclear Factor Erythroid 2-related Factor 2 (NRF2), Independent of Transcription Factor TFEB. J Biol Chem. 2015;290(24):14945–62.

58. East DA, Fagiani F, Crosby J, Georgakopoulos ND, Bertrand H, Schaap M, et al. PMI: a DeltaPsim independent pharmacological regulator of mitophagy. Chem Biol. 2014;21(11):1585–96.

59. Gumeni S, Papanagnou ED, Manola MS, Trougakos IP. Nrf2 activation induces mitophagy and reverses Parkin/Pink1 knock down-mediated neuronal and muscle degeneration phenotypes. Cell Death Dis. 2021;12(7):671.

60. Sarlette A, Krampfl K, Grothe C, Neuhoff N, Dengler R, Petri S. Nuclear erythroid 2-related factor 2-antioxidative response element signaling pathway in motor cortex and spinal cord in amyotrophic lateral sclerosis. J Neuropathol Exp Neurol. 2008;67(11):1055–62.

61. Nardo G, Iennaco R, Fusi N, Heath PR, Marino M, Trolese MC, et al. Transcriptomic indices of fast and slow disease progression in two mouse models of amyotrophic lateral sclerosis. Brain. 2013;136(Pt 11):3305–32.

62. Moujalled D, Grubman A, Acevedo K, Yang S, Ke YD, Moujalled DM, et al. TDP-43 mutations causing amyotrophic lateral sclerosis are associated with altered expression of RNA-binding protein hnRNP K and affect the Nrf2 antioxidant pathway. Hum Mol Genet. 2017;26(9):1732–46.

63. Jimenez-Villegas J, Kirby J, Mata A, Cadenas S, Turner MR, Malaspina A, et al. Dipeptide Repeat Pathology in C9orf72-ALS Is Associated with Redox, Mitochondrial and NRF2 Pathway Imbalance. Antioxidants (Basel). 2022;11(10).

64. Vucic S, Henderson RD, Mathers S, Needham M, Schultz D, Kiernan MC, et al. Safety and efficacy of dimethyl fumarate in ALS: randomised controlled study. Ann Clin Transl Neurol. 2021;8(10):1991–9.

